# Reduced mitochondria provide an essential function for the cytosolic methionine cycle

**DOI:** 10.1101/2022.04.01.486701

**Authors:** Justyna Zítek, Zoltán Füssy, Sebastian C. Treitli, Priscila Peña-Diaz, Zuzana Vaitová, Daryna Zavadska, Karel Harant, Vladimír Hampl

## Abstract

It has been long hypothesised that mitochondrial reduction is intrinsically related to the remodelling of Fe-S clusters assembly. Yet as our knowledge of divergent free-living protists broadens, so does the spectrum of variability within the range of mitochondrial-related organelles (MROs) fundamental functions. We resolved to high precision the MRO proteome of *Paratrimastix pyriformis* using Localisation of Organelle Proteins by Isotope Tagging (LOPIT) and demonstrate its role in the synthesis of folate derivates bearing one-carbon (1C) units, its link to the glycine cleavage system (GCS) and their only conceivable role as suppliers for the cytosolic methionine cycle, involved in recycling of S-adenosine methionine. This observation provides congruity to the presence of GCS in MROs of free-living anaerobes and its absence in endobionts, which typically lose the methionine cycle and, in the case of oxymonads, also mitochondria.

## Introduction

It is now well-established that the mitochondrion has been acquired prior to the last eukaryotic common ancestor (LECA) and is present in all eukaryotes with the single known exception of oxymonad *Monocercomonoides exilis* (Karnkowska et al., 2016; Roger et al., 2017). Eukaryotes thriving in low-oxygen environments usually display divergent, reduced types of mitochondria (mitochondrion-related organelles, MROs), the best-studied examples of which are hydrogenosomes and mitosomes (Santos et al., 2018). The diversity of MROs in free-living protists has been revealed recently (Leger et al., 2017), but functional methodologies are hindered by their xenic cultivation methods. Their organelles are characterised using transcriptomic and/or genomic data, targeting predictions, and homology searches that use established sets of MRO proteins as baits. Though this methodology has proven very powerful it suffers from serious biases. For instance, the error rates of targeting predictors are typically unknown and are expected to vary considerably between organisms. Homology-based methods, in contrast, cannot recover proteins that are not already in the researcher’s crosshairs. Moreover, the assumption that orthology translates into identical cellular localisation is not entirely correct. As a result, heterologous localisation in tractable model organisms endure the hardship of inaccuracy and their validity roughly corresponds to the similarity between the tested organelle and its heterologous model.

In view of all mentioned shortcomings, experimental, self-standing *de novo* identifications of MRO proteomes by proteomic methods are urgently/gravely required, as such data would radically enhance *in-silico* predictions of MRO proteomes, yet their availability is rather scarce. The few published proteomes of reduced mitochondria belong to only three lineages of parasitic or endocommensal protists, namely the hydrogenosomes of trichomonads, the hydrogenosomes or mitosomes of diplomonads and the mitosome of *Entamoeba histolytica* (Beltrán et al., 2013; Fang et al., 2016; Jedelský et al., 2011; Jerlström-Hultqvist et al., 2013; Mi-ichi et al., 2009; Rada et al., 2011; Schneider et al., 2011).

To the best of our knowledge, the oxymonad flagellate Monocercomonoides exilis (Preaxostyla, Metamonada) is completely devoid of mitochondria and as such it represents a notable exception amongst eukaryotes. Evidence suggests that in the common ancestor of Preaxostyla the mitochondrial pathway for iron-sulfur cluster assembly (ISC) had been replaced by the sulfur mobilisation system (SUF), acquired by lateral gene transfer from unknown bacteria (Karnkowska et al., 2016; Vacek et al., 2018). This replacement stripped the MRO of one of its essential functions (Braymer et al., 2021), hence preadapting the lineage to its subsequent loss. Regardless, not all members of Preaxostyla are amitochondriate, thus the correlation between the presence of the SUF pathway and the absence of mitochondrion is not rigorous and MROs likely perform other essential function(s) in species that retained them. *Paratrimastix pyriformis*, a free-living freshwater bacterivorous flagellate, is one such case and its MRO has been partially characterised (Hampl et al., 2008; O’Kelly et al., 1999; Zubáčová et al., 2013). A precise determination of the protein composition and metabolic roles of the MRO in *P. pyriformis* is thus key to understanding the process of MRO reduction and its loss in Preaxostyla.

The only experimentally localised pathway in this organelle is the glycine cleavage system (GCS, composed of four subunits H, L, P, and T), involved in the degradation of glycine (Zubáčová et al., 2013). GCS-H was localised *in situ* by immunofluorescence staining, while the localisation of GCS-P was inferred using heterologous systems. Many other crucial questions regarding the biochemical function of this MRO remain unanswered, namely the localisation of Fe-S cluster assembly and that of extended glycolysis enzymes. The latter represents a common repertoire of proteins present in anaerobic and microaerophilic protists forming acetyl-CoA and providing a sink for electrons from reduced ferredoxins and NADH (Hrdy et al., 2004) exemplified by pyruvate:ferredoxin oxidoreductase (PFO) and [FeFe] hydrogenase (HydA), respectively. In the transcriptome of *P. pyriformis*, several genes encoding HydA and PFO are present, as well as three maturases (HydE, HydF and HydG) (Hampl et al., 2008; Zubáčová et al., 2013) essential for the assembly of an active site of HydA (McGlynn et al., 2007). Although some of the above proteins bear N-terminal targeting extensions, experimental evidence for their MRO localisation is missing (Zubáčová et al., 2013).

We reconstructed the functions of *P. pyriformis* MRO using spatial proteomic data. The established MRO proteome contains 31 proteins including a complete pathway of folate-mediated one-carbon (1C) metabolism, a route connected to GCS and capable of producing formate. We propose that the production of formate and methylated folate species remains an unnoticed yet essential function of MROs in free-living eukaryotes.

## Results

### Applying LOPIT-DC robustly assigned more than a thousand proteins into sub-compartments of *P. pyriformis* cell

We applied Localisation of Organelle Proteins by Isotope Tagging after Differential UltraCentrifugation (LOPIT-DC) to determine proteins localisation within *P. pyriformis* cells (Geladaki et al., 2019). The method relies on the assumption that proteins localised in the same organelle co-fractionate, hence displaying similar distribution profiles to known organellar markers (Sadowski et al., 2006). Cell lysates were divided by differential centrifugation into 10 fractions – 9 fractions of pelleted cell components and supernatant enriched in the remaining cytosolic proteins (Figure 1A). Inspection of fractions 1-6 using transmission electron microscopy (TEM) ensured that the protocol did not compromise the integrity of organellar membranes (Figure S1) and confirmed conspicuous changes of fraction content with MROs present mainly in fractions #2 and #3.

**Figure 1.**
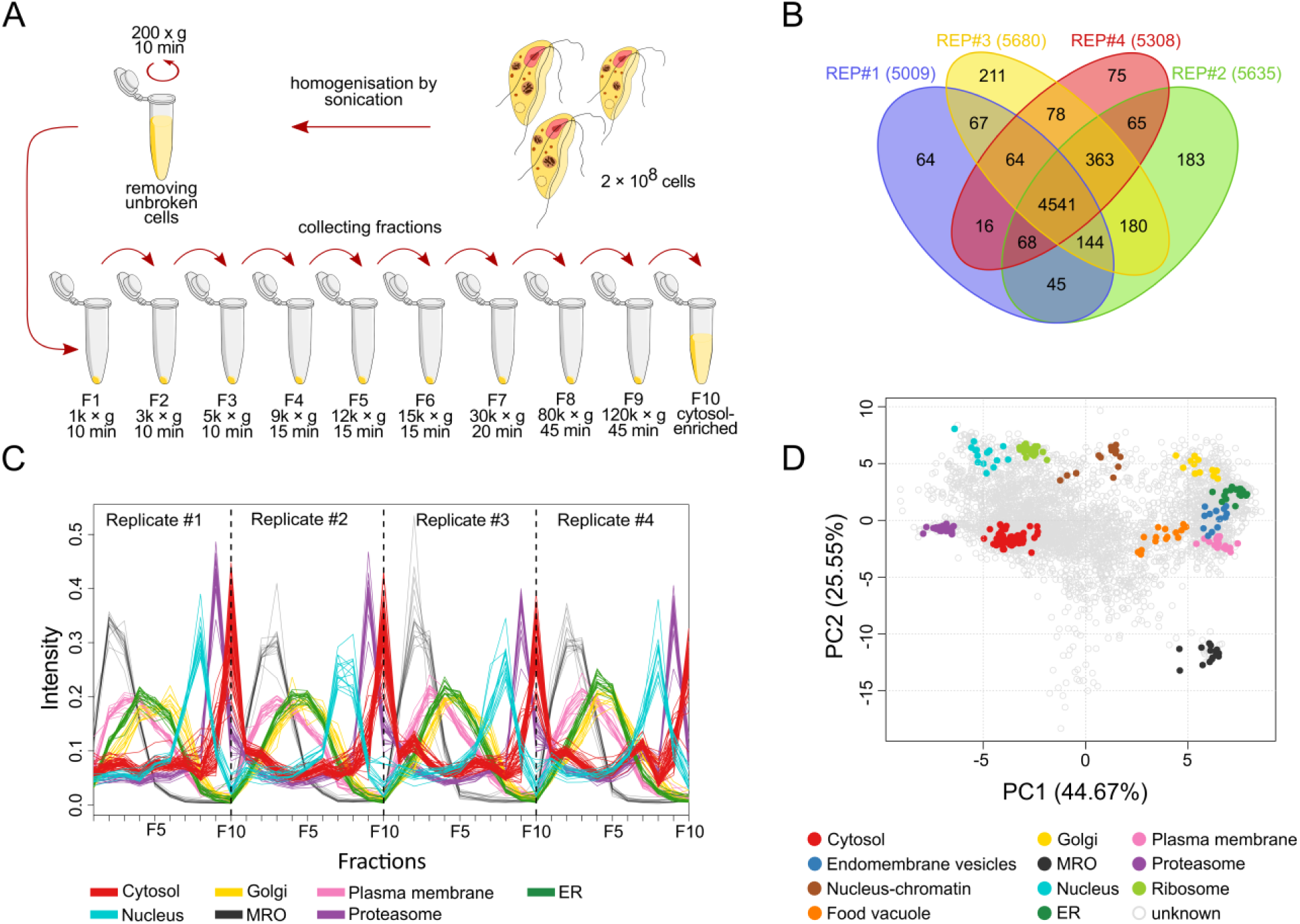
Application of LOPIT-DC on P. pyriformis cells. A) Schematic representation of the fractionation workflow. B) Venn diagram displaying common and unique proteins identified in four replicates. The total number of quantified proteins for each replicate is shown in brackets. C) Distribution profile of marker proteins across ten labelled fractions, for clarity only seven selected sub-compartments are shown. D) Principal component analysis (PCA) plot mapping 226 markers proteins for 11 sub-compartments.

Four biological replicates of similar sets of 10 fractions were prepared, labelled with tandem mass tag (TMT) 10-plex reagent, and peptide abundances across them were quantified by mass spectrometry. More than 5,000 proteins were identified in each biological replicate, 4,541 of which were common for all (Figure 1B, Table S1) and this common subset was further analysed in R using pRoloc package (Gatto et al., 2014).

Classification using supervised machine learning methods requires a manually curated set of markers, a list of proteins of well-known localisation. For the purpose of this analysis, we compiled a set of 226 markers for 11 sub-compartments (Table S2) based on up-to-date manually curated annotations, data from previous studies (Zubáčová et al., 2013) and unsupervised or semi-supervised approaches (see Methods for details). The distribution profiles of marker proteins for 7 selected sub-compartments displayed high similarities between replicates, suggesting acceptable reproducibility of the fractionation protocol (Figure 1C). The distribution profile of nuclear proteins, or lack thereof in the first fraction, indicates that either the nuclei were removed together with unbroken cells or disrupted by sonication. The MRO exhibited the most distinguished profile amongst all sub-compartments, and it was enriched mainly in fractions #2 and #3, corresponding to EM observations (Figure S1).

Visualisation of protein distribution in a 2D plot was achieved by principal component analysis (PCA). Marker proteins were mapped on the PCA projection (Figure 1D) and amongst well-separated sub-compartments we could clearly distinguish the components of the MRO. To quantify the resolution between sub-compartments, we used the Qsep score (Gatto et al., 2019) which provided ratios of between-cluster to within-cluster average distances (Figure S2). This analysis confirmed that the MRO is well-defined and separated from other sub-compartments.

To assign the remaining 4,315 proteins into sub-compartments we used two classification algorithms: support vector machine (SVM) and Bayesian approach based on t-augmented Gaussian mixture model with *maximum a posteriori* prediction (TAGM MAP) (Figure 2). In total, 2,503 proteins were classified into one of the sub-compartments by at least one algorithm, and in 1,012 cases both algorithms agreed; 1,786 proteins remained without classification (Table S2). In general, SVM classified higher number of proteins overall, particularly of the cytosolic sub-compartment, while TAGM MAP tended to inflate the nuclear sub-compartment. The MRO and ribosome sub-compartments displayed the most consistent results between both methods.

**Figure 2.**
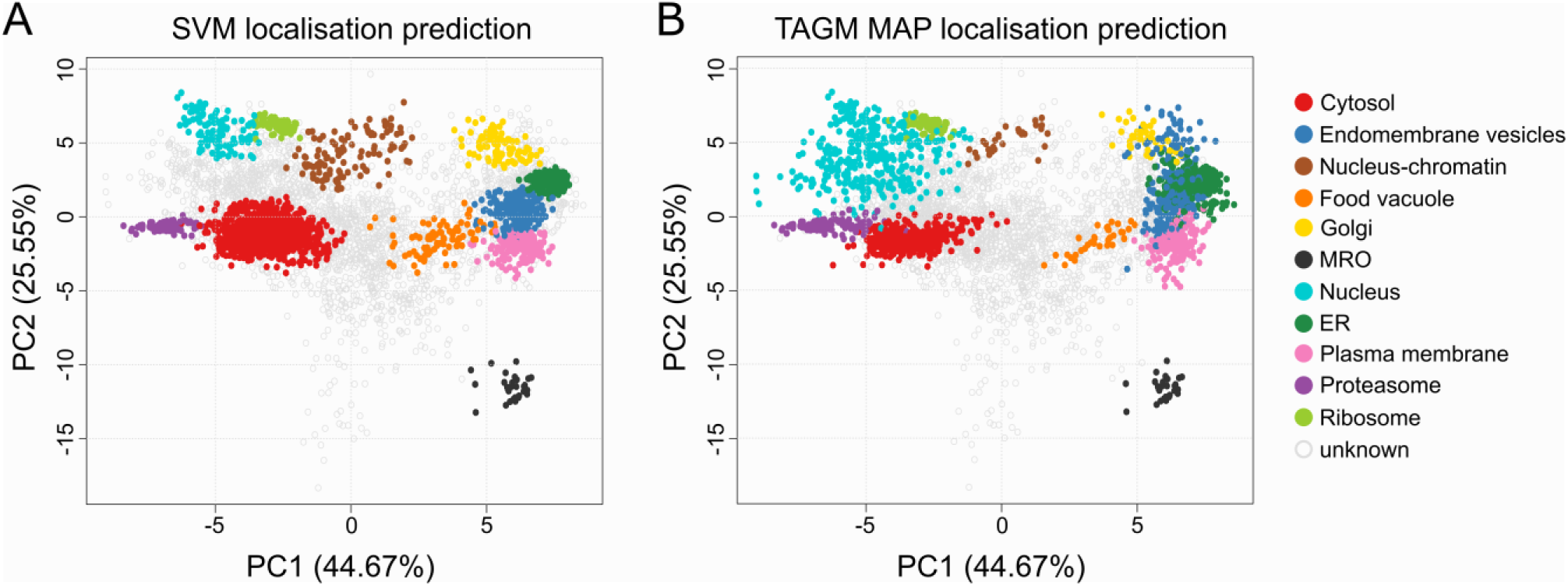
PCA plots with proteins assigned to sub-compartments labelled. A) PCA projection after SVM classification B) PCA projection after TAGM MAP classification

### The mitochondrion-related organelle provides THF intermediates to cytosolic pathways

Altogether, 31 proteins (including 16 markers) were assigned to the MRO (Table 1), and of the 15 proteins assigned *de novo*, both classification methods agreed in 13 cases. The protein composition of the MRO was thus resolved with high confidence.

**Table 1.**
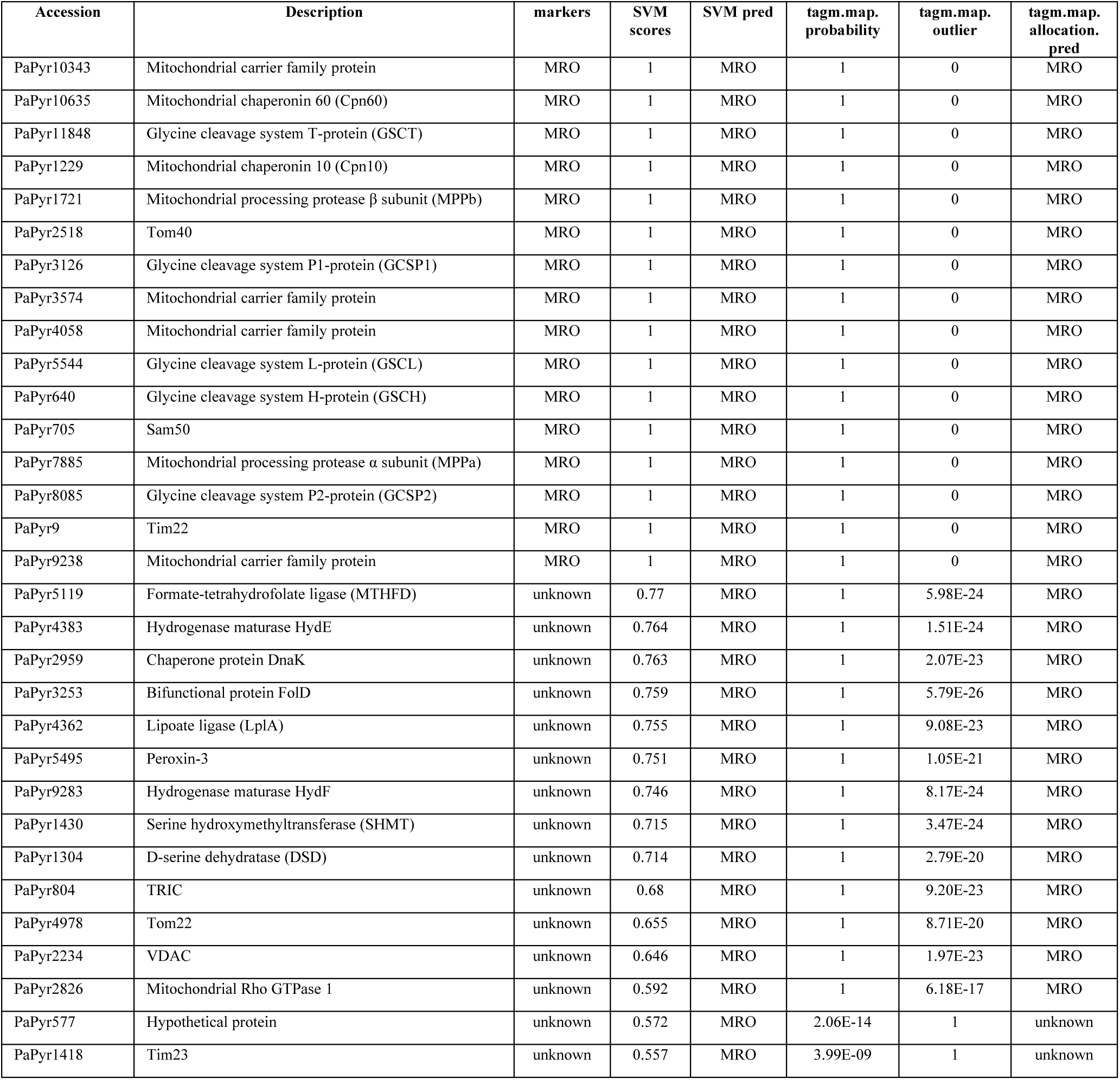
P. pyriformis MRO proteome established by LOPIT-DC analysis. List of proteins assigned to the MRO compartment. The list includes 16 marker proteins and 15 proteins identified in this study. Scores and predictions by two statistical methods are provided.

The map of organelle functions built on predicted protein content is shown in Figure 3. Members of the mitochondrial carrier family (MCF), proteins involved in the transport and maturation of MRO proteins, and components of the GCS were used as markers. Amongst the *de novo* assigned MRO proteins were bifunctional methylenetetrahydrofolate dehydrogenase/methenyl tetrahydrofolate cyclohydrolase (FolD; PaPyr3253), formate-tetrahydrofolate ligase (MTHFD; PaPyr5119) and serine hydroxymethyltransferase (SHMT; PaPyr1430). These three proteins play a role in folate-mediated 1C metabolism where formate is generated from serine. The first reaction catalysed by SHMT (PaPyr1430) brings a 1C unit into the folate pool by transferring a methyl group from serine to tetrahydrofolate (THF). Indeed, PaPyr1430 has been previously considered an MRO protein based on its N-terminal extension (Zubáčová et al., 2013). Further interconversion of THF derivates into formate is mediated by FolD (PaPyr3253) and MTHFD (PaPyr5119), leading to the concurrent production of ATP (Figure 3). Finally, folylpolyglutamate synthase (FPGS; PaPyr1465), an enzyme important for intracellular folates retention and compartmentalisation (Lawrence et al., 2014), seems to be dually localised in both the MRO and cytosol, judging from its mixed distribution profile (Figure S3B).

**Figure 3.**
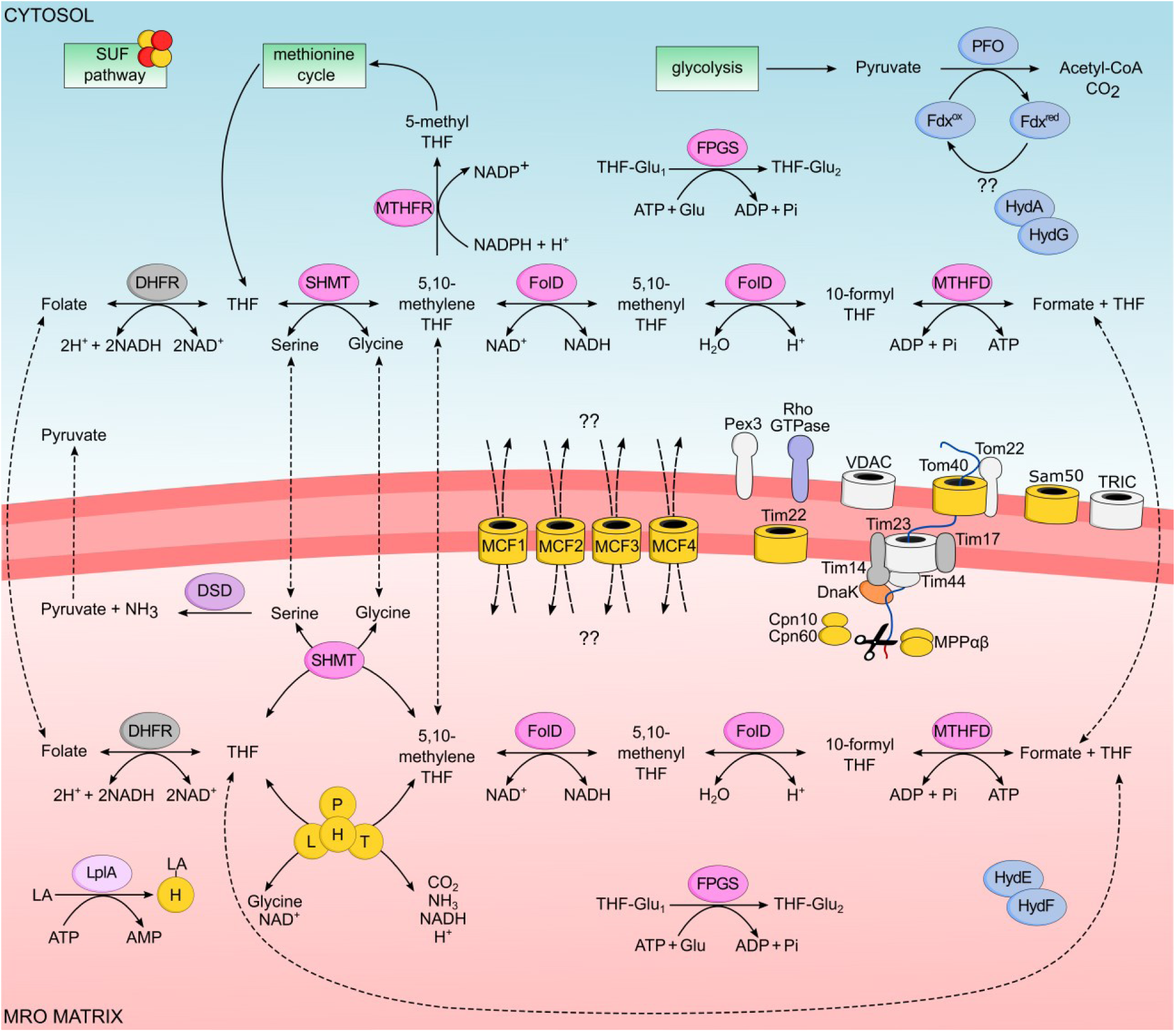
Metabolic map of P. pyriformis MRO based on LOPIT-DC analysis. Proteins used as markers for classification are coloured in yellow, hypothetical proteins are coloured grey, proteins present in the genome but missing in LOPIT dataset are coloured dark grey. H, L, P, T – components of glycine cleavage system; SHMT, serine hydroxymethyltransferase; FolD, methylenetetrahydrofolate dehydrogenase/methenyl tetrahydrofolate cyclohydrolase; MTHFD, formate-tetrahydrofolate ligase; FPGS, folylpolyglutamate synthase; MTHFR, methylenetetrahydrofolate reductase; DHFR, dihydrofolate reductase; LplA, lipoate ligase; DSD, D-serine dehydratase; PFO, pyruvate:ferredoxin oxidoreductase; Fdx, ferredoxin; HydA, [FeFe] hydrogenase; HydE, HydF, HydG, hydrogenase maturases; MPPαβ, mitochondrial processing protease subunits α and β; VDAC, voltage-dependent anion channel; TRIC, trimeric intracellular ion channel; Pex3, peroxisomal biogenesis factor 3; Cpn10, mitochondrial chaperonin 10; Cpn60, mitochondrial chaperonin 60; DnaK, chaperone protein; MCF1, MCF2, MCF3, MCF4, mitochondrial carrier family proteins; THF, tetrahydrofolate; LA, lipoic acid.

Folate-mediated 1C metabolism has been mainly studied in human cancer cells, considered a potential target for treatment due to its up-regulation (Nilsson et al., 2014). In mammalian cells, the complete 1C pathway is localised both in the cytosol and the mitochondria (Tibbetts and Appling, 2010). This situation was observed also in *P. pyriformis*, where the second paralogues for each, MTHFD (PaPyr5307), FolD (PaPyr2026) and SHMT (PaPyr5925), were classified clearly as cytosolic (Figure 3, Table S2) together with another enzyme related to 1C metabolism, methylenetetrahydrofolate reductase (MTHFR; PaPyr5097). MTHFR catalyses the irreversible conversion of 5,10-methylene-THF into 5-methyl-THF, substrate of the methionine cycle (Mudd et al., 2007). Considering all this, we hypothesise that formate or THF species produced within the MRO are exported to the cytosol, where they are utilised for production of 5-methyl-THF supplementing the methionine cycle. All enzymes of the methionine cycle were detected and classified as cytosolic.

The remaining proteins classified as MRO-localised were lipoate ligase (LplA, PaPyr4362), Rho GTPase (PaPyr2826), chaperone protein DnaK (PaPyr2959) and D-serine dehydratase (DSD, PaPyr1304). Lipoic acid (LA) is an essential cofactor in subunits of four known enzyme complexes, all of them generally present in mitochondria (Rowland et al., 2018), yet only GCS-H is present in *P. pyriformis*. Lipoylation of H-protein is likely mediated by LplA (PaPyr4362) that plays role in a bacterial salvage pathway, where exogenous lipoic acid is activated and transferred to H-protein (Green et al., 1995; Solmonson and DeBerardinis, 2018). Mitochondrial Rho GTPases, known also as Miro, localise in the outer membrane and contain two Ca^2+^ binding EF-hands domains flanked by GTPase domains on both sides (Fransson et al., 2003). Studies on various models suggested that Miro proteins are important for mitochondrial transport within the cell and maintenance of their morphology and homeostasis (Fransson et al., 2003; Saotome et al., 2008). *P. pyriformis* Rho GTPase (PaPyr2826) is shorter than Miro proteins from other eukaryotes as it lacks a GTPase domain at the C-terminus. D-serine dehydratase (PaPyr1304) is a PLP-dependent enzyme converting serine into pyruvate and ammonia (Dowhan and Snell, 1970). Chaperone protein DnaK (PaPyr2959) is a proteobacterial ortholog of mitochondrial Hsp70, a pivotal player in mitochondrial protein import (Boorstein et al., 1994).

### The dark proteome of the MRO likely consists of membrane proteins involved in its biogenesis

Six proteins robustly assigned to the MRO (PaPyr577, PaPyr804, PaPyr1418, PaPyr2234, PaPyr4978, PaPyr5495) lack functional annotation and thus have been labelled as hypothetical. To mine information regarding the identity/function of these proteins, we searched for conserved motifs and transmembrane domains and used more sensitive profiling and structure-informed methods [HHpred (Söding, 2005), HMMER v3.3, hmmer.org, SWISS-MODEL (Bienert et al., 2017)] to find putative homologues in other organisms. Using the same tools, we examined yet another hypothetical protein, PaPyr7077, which regardless of lacking MRO assignation by machine learning approaches exhibited a distribution profile consistent with MRO proteins in two sample replicates. We propose functions for six of them.

PaPyr2234 is a very abundant protein in our dataset, and it is likely a divergent member of the porin3 superfamily (PF01459), potentially representing a voltage-dependent anion channel (VDAC) according to SWISS-MODEL. VDACs localise in the outer membrane of mitochondria and are important for the passive diffusion of ions and small metabolites, including ATP/ADP, NAD/NADH, pyruvate, glutamate, or malate (Gincel and Shoshan-Barmatz, 2004; Rostovtseva and Colombini, 1997). PaPyr804 represents according to HHpred search and phylogenetic reconstruction (Figure S4) a trimeric intracellular ion channel (TRIC). In eukaryotes, these monovalent cation channels are predominantly present in the membrane of the endoplasmic reticulum (ER) where they balance the charge during Ca^2+^ uptake or release (Yazawa et al., 2007).

PaPyr4978, PaPyr7077, and PaPyr1418 were recognised as members of the TOM/TIM translocase machinery, Tom22, Tim44, and Tim17/22/23, respectively. It has been shown in yeast that three different proteins (Tom20, Tom70 and Tom22) help recognise proteins transported into mitochondria (Brix et al., 1997). In the genome of *Paratrimastix pyriformis*, homologues of Tom20 and Tom70 failed to be identified, hence PaPyr4978 is putatively the one to recognise targeting peptides on proteins directed to the MRO. PaPyr7077 represents a putative homologue of Tim44, a component of the PAM motor complex (Neupert and Brunner, 2002). The presence of Tim44 has been reported in other Metamonads (Beltrán et al., 2013; Leger et al., 2017; Pyrihová et al., 2018; Rada et al., 2011; Schneider et al., 2011; Stairs et al., 2021) and our phylogenetic reconstruction pinpoints the relationship of PaPyr7077 to Tim44 of *Trichomonas vaginalis* (Figure S4). Members of the Tim17/22/23 protein family have been identified in all eukaryotes as components of protein translocation complexes in the mitochondrial inner membrane – TIM23 complex facilitates protein import into the matrix, while TIM22 mediates protein insertion into the mitochondrial inner membrane (Chaudhuri et al., 2020; Rassow et al., 1999; Žárský and Doležal, 2016). In the genome of *P. pyriformis*, two other obvious homologues of Tim17/22/23 proteins are present (PaPyr9 and PaPyr5590), of which the former has been detected here and used as a marker. Both contain the conserved glycine-zipper (GX_3_GX_3_G) and PRAT ((G/A)X_2_(F/Y)X_10_RX_3_DX_6_(G/A/S)GX_3_G) motifs, hallmarks of this protein family. In PaPyr1418 only a glycine-zipper motif was identified. Phylogenetic affiliations of *P. pyriformis* Tim17/22/23 proteins were determined using the dataset of Žárský and Doležal (Žárský and Doležal, 2016) enriched by sequences from diverse eukaryotic lineages not available in the Uniprot database (Richter et al., 2020), resulting in PaPyr5590, PaPyr9, and PaPyr1418 as orthologues of Tim17, Tim22, and Tim23, respectively (Figure 4).

**Figure 4.**
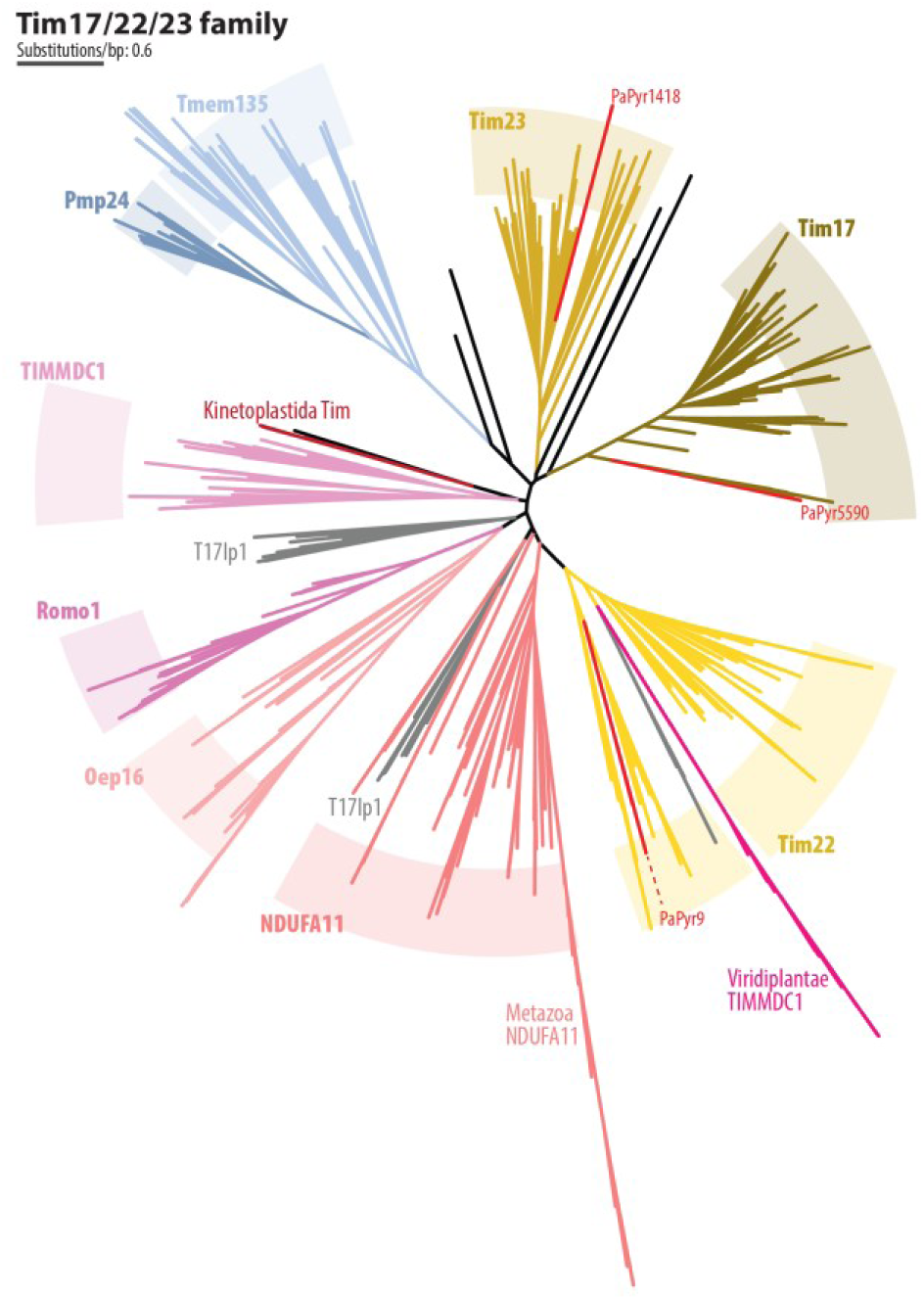
Schematic phylogeny of the Tic17/22/23 family with the positions of P. pyriformis homologs. Employing a previously compiled dataset (Žárský and Doležal, 2016), the ML tree was inferred by IQ-TREE using the Posterior Mean Site Frequency empirical model with the ultrafast bootstrapping strategy (see Methods).

Lastly, a putative function was assigned to PaPyr5495, a protein sharing similarities with peroxisomal biogenesis factor 3 (Pex3) (Figure S4). In human cells, Pex3 is responsible for docking of receptor protein Pex19 on the peroxisomal membrane during the import of PTS1 C-terminal signal-containing proteins (Fang et al., 2004). Importantly, Pex19 homologue in *P. pyriformis* (PaPyr8690) was identified using HMM profiles and shown to be phylogenetically related to other sequences from Preaxostyla (Figure S4). All Preaxostyla Pex19 homologues clearly possess a conserved Pex19 Pfam motif (PF04614) and, yet divergent, *Trimastix marina* and *P. pyriformis* Pex19 bear an N-terminal sequence similar to the Pex3-interaction motif known from plants, fungi, and animals (Figure S4). Since no peroxisomes have been observed in Preaxostyla (Hampl, 2017) and no other genetic markers for peroxisomes were detected (Karnkowska et al., 2016, 2019) we speculate that the Pex3/Pex19 system may have been recruited to function in the MRO membrane of *Paratrimastix pyriformis*.

### The MRO is not involved in extended glycolysis and Fe-S clusters assembly

In the hydrogenosomes of trichomonads, pyruvate undergoes oxidative decarboxylation into acetyl-CoA and CO_2_ by PFO, while ferredoxins reduced in this reaction are subsequently reoxidized by HydA with concomitant production of H_2_ (Lindmark et al., 1975). Five genes encoding for PFO, two for ferredoxins, nine for HydA, and one gene for each of the three HydA maturases (HydE, F, G) are present in the *P. pyriformis* genome.

Three out of four PFOs (with the sole exception was PFO_3, PaPyr7655) were identified by LOPIT-DC and assigned to the cytosolic compartment by at least one classification method. Both genes annotated as ferredoxin (PaPyr13543, PaPyr13544) showed a distribution profile similar to cytosolic markers but were not classified as cytosolic by algorithms (Figure S3A.). Five out of nine HydA enzymes were detected in all four LOPIT-DC replicates and none of them exhibited a distribution profile that would suggest localisation into the MRO (Figure S3A). HydA_4 (PaPyr6388) was assigned to the cytosol by both classification tools, whereas HydA_1 (PaPyr1254), HydA_2 (PaPyr1278) and HydA_6 (PaPyr8158) were assigned to the cytosol only by SVM. The last hydrogenase present in the dataset, HydA_8 (PaPyr8532_PaPyr8533), was not classified to any sub-compartment. Of hydrogenase maturases, curiously, two (HydE; PaPyr4383 and HydF, PaPyr9283) were robustly assigned to the MRO while the profile of the third (HydG, PaPyr1252) was clearly cytosolic (Figure S3A). Altogether, these results support a cytosolic localisation of extended glycolysis in *P. pyriformis* but raise an interesting question on the role of hydrogenase maturases.

The common function for virtually all mitochondria and MROs is the synthesis of Fe-S clusters, in Preaxostyla performed by the sulphur mobilisation system (SUF). This pathway consists of the ATPase SufC (PaPyr9323), a scaffold protein SufB (PaPyr903) and the cysteine desulfurase fusion protein SufDSU (PaPyr904) (Vacek et al., 2018). All three proteins were detected in all four LOPIT-DC replicates and enriched in the cytosolic fraction (Figure S3A). Both classification methods assigned SufC and SufDSU to the cytosol; SufB was classified as cytosolic only by the SVM algorithm, but with high confidence. The data clearly point to cytosolic localisation of Fe-S cluster synthesis in *P. pyriformis*.

### Localisation by heterologous expression often contradicts the results of LOPIT-DC

A popular experimental approach for assessing the localisation of proteins from organisms that cannot be genetically transformed is the expression in heterologous systems. *T. vaginalis* and its hydrogenosome are evolutionarily related to the MRO of *P. pyriformis*, therefore we have chosen this model to attest to the localisation of 14 selected non-marker proteins classified robustly to the MRO or the cytosol. C-terminally HA-tagged proteins were expressed in *T. vaginalis* and their localisation was visualised by immunofluorescence microscopy (IFA) and compared with an anti-malic enzyme antibody used as hydrogenosomal marker (Figure S5). Of all tested proteins, 5 colocalised with malic enzyme staining, HydE (PaPyr4383), HydA_4 (PaPyr6388), HydA_6 (PaPyr8156) putative VDAC (PaPyr2234), putative Pex3 (PaPyr5495); 7 proteins displayed cytosolic localisation, HydA_2 (PaPyr1278), HydG (PaPyr1252), PFO_2 (PaPyr12211), PFO_4 (PaPyr7937_PaPyr7938), FolD (PaPyr3253), MTHFD (PaPyr5119) and SufC (PaPyr9323). The last two proteins, HydF (PaPyr9283) and putative Tom22 (PaPyr4978) exhibited patterns that could not be safely interpreted either way. In many cases (HydA_2, HydG, both PFOs, FolD, MTHFD, SufC, VDAC) the hydrogenosomal signal appeared unusual suggesting that the overexpression may cause a phenotype that affects hydrogenosomes. These results should therefore be carefully interpreted.

In summary, the results of heterologous expression localisations matched the outcomes of LOPIT proteomics in only 8 cases and were often hard to interpret. As the sub-compartmental classification of the tested proteins by LOPIT-DC was robust and difficult to dispute, the result indicates that the heterologous approach, at least in the case of *P. pyriformis*, is of little suitability due to high artifactuality.

### N termini of MRO proteins have a low positive charge and are not reliably predicted as targeting signals

The translation of most proteins destined to aerobic mitochondria takes place in the cytosol and their delivery is ensured by N-terminal or internal targeting sequences (Wiedemann and Pfanner, 2017). Mitochondrial N-terminal targeting signals (NTS) vary in length, but they share some common features – they are cleaved by mitochondrial processing peptidases (MPP) within an arginine-containing motif and display an amphipathic α-helix structure composed of hydrophobic and positively charged amino acids (Kunze and Berger, 2015; Roise et al., 1986). Bioinformatic tools that predict mitochondrial localisation largely focus on these N-terminal features, but in proteins targeted to MROs, divergence hinders their identification. Meanwhile, internal localisation signals cannot be predicted at all (Mentel et al., 2008; Zimorski et al., 2013). We have used our experimentally determined set of MRO proteins of *P. pyriformis* to calculate the average net charge of their N-termini and tested the performance of targeting predictors for this organism.

The average net charge of the first 10, 20 and 30 amino acids of all 31 MRO proteins was calculated for three pH values using Isoelectric Point Calculator 2.0 (http://ipc2.mimuw.edu.pl/index.html; Table S3). At neutral pH, the N-termini of most proteins were calculated to contain a weak positive net charge close to 1.0 and only in two proteins, mitochondrial membrane carrier 4 (PaPyr9238) and Pex3 (PaPyr5495), the value was above 3.0. Values are similar to those in *T. vaginalis* hydrogenosomal proteins but lower than those in yeast, human, or *Anaeramoeba*, where they fall in the range 3.0-4.0 (Garg and Gould, 2016; Garg et al., 2015; Stairs et al., 2021). This suggests that the positive charge of N-termini has low importance for the delivery of proteins into *P. pyriformis* MRO and that this MRO likely displays weak or no membrane potential.

The performance of 6 tools for mitochondrial targeting prediction (iPSORT, MitoFates, NommPred, MultiLoc2, TargetP 2.0 and Cello) was tested using the set of MRO proteins and 10 negative control sets of the same sizes (n=30) robustly assigned to 10 other sub-compartment using LOPIT-DC (Table S4). Specificity was always below 50% (the highest value 0.6 by TargetP-2.0, which however displayed poor sensitivity of 0.17), i.e., more than half of the predicted proteins were false positives. The highest measured sensitivity was 0.63 (by Cello; 0.82 on the soluble mitochondrial subset). Although the algorithms tend to select MRO proteins from the set, their error rates were still too high to be useful for genome-wide prediction of *P. pyriformis* MRO proteins.

## Discussion

The hyperplexed LOPIT (hyperLOPIT) approach was shown to be a powerful method to map protein distribution within cells (Christoforou et al., 2016) and its recent application to *Toxoplasma gondii* tachyzoites proved successful also for protists (Barylyuk et al., 2020). Yet, hyperLOPIT is not suitable for all cell types due to a large amount of starting material required to perform sub-cellular fractionation by density gradient (Mulvey et al., 2017). For polyxenically grown *P. pyriformis* with low cell densities, we employed a simplified version – LOPIT-DC, which produces comparable results with less starting material (Geladaki et al., 2019). 4,541 *P. pyriformis* proteins showed measurable abundance profiles in all four biological replicates. This number is comparable to the number of proteins detected in human U-2 OS cells (6,837 proteins), considering the total size of predicted proteomes (Geladaki et al., 2019), as well as the fraction of proteins successfully classified into sub-compartments. Geladaki et al. (2019) classified 35% of identified proteins using SVM method, while 45% were classified by the same method in this study. Nevertheless, it should be noted that the number of unclassified proteins depends on factors such as quality of the marker set, threshold setup and dynamics of protein localisations within the cell, which if high may complicate the classifications (Thul et al., 2017). Finally, the median distance between the clusters calculated by Qsep in our experiment is similar to that of Geladaki et al. (2019), suggesting that we have reached a satisfactory level of clusters separation.

To analyse LOPIT data by machine learning methods, it is recommended to use at least 13 markers per sub-compartment (Breckels et al., 2018). For model organisms, e.g., *Homo sapiens*, a marker set of 872 proteins for 12 sub-compartments is available in the pRoloc package (Breckels et al., 2018). Due to the virtual absence of experimental data, our marker set for 11 sub-compartments comprised only 226 proteins, hence the predicted clusters of proteins may be less accurate than in human cells or *T. gondii*. Therefore, we decided to focus only on the proteome of the MRO for which we used 16 reliable markers proteins and displayed the best separation from other sub-compartments.

Thirty-one proteins assigned to the MRO sub-compartment is a relatively small number compared to the proteome of *T. vaginalis* hydrogenosome that comprises potentially as many as 600 proteins (Beltrán et al., 2013; Rada et al., 2011; Schneider et al., 2011). Given that we experimentally detected only about one-third of the total predicted proteome, we hypothesise that the MRO proteome is not complete, as some proteins escaped the detection due to the sensitivity limit or were not expressed under experimental conditions. Even considering these limitations, LOPIT data provided us with the most reliable picture of protein localisation, less prone to contaminations than classical proteomics based on organelle purification. Notably, Beltrán et al. (2013) identified 631 proteins in the fraction of highly purified *T. vaginalis* hydrogenosomes, but only 179 were considered true hydrogenosomal proteins and out of 2,934 proteins identified in the hydrogenosome of *Pentatrichomonas hominis*, only 442 were considered reliable candidates (Fang et al., 2016).

The results of the LOPIT analysis revealed the presence of a complete folate-mediated 1C metabolism in the *P. pyriformis* MRO, extending the previously experimentally localised GCS. The GCS and the folate pathway in eukaryotes are intrinsically interconnected to deliver 1C units for methylation, methionine regeneration, and *de novo* formation of purines and thymidine through various folate species which serve as 1C carriers (Ducker and Rabinowitz, 2017). In mammals, compartmentalisation of the 1C pathway between cytosol and mitochondria is thought to occur to avoid depletion of cytosolic NAD^+^ required for glycolysis, therefore oxidative direction of the reactions is favoured in mitochondria while reductive in the cytosol (Tibbetts and Appling, 2010). *P. pyriformis* lacks most routes that require different forms of THF, hence 1C metabolism likely plays one single role, namely the synthesis of 5-methyl-THF, essential for the methionine cycle. In this cycle, methylation of homocysteine regenerates methionine, which is further converted to S-adenosyl methionine (SAM), an essential cofactor for a great number of reactions (Mudd et al., 2007).

Whilst reactions of 1C metabolism are reversible, their reversibility depends on the availability of substrates and their stoichiometric amounts. This premise is observed as THF upon the synthesis of formate by MTHFD, is released and ready to cycle with a newly catabolised carbon unit produced by the activity of either SHMT or GCS, within the organelle. Hence it is accepted that the carbons from 1C metabolism are required in stoichiometric amounts, meanwhile, THF is only required in catalytic amounts (Tibbetts and Appling, 2010). In *P. pyriformis*, in absence of substrate-level or oxidative phosphorylation, the requirement of the MRO to replenish NAD^+^ used by FolD and GCS likely depends on the activity of DHFR, a step that requires folate entry in the organelle with concomitant oxidation of 2 molecules of NADH for every carbon that enters the GCS, which would force a stricter 1C:THF stoichiometry in the organelle. Two genes encoding DHFR are present in the genome of *P. pyriformis* (PaPyr1795, PaPyr350), neither of which was detected in the LOPIT-DC analysis. The identity of 1C donors in *P. pyriformis* can only be inferred from the enzymes found in the organelle for their catabolism, indicating that the donor molecules are likely serine, glycine and formate. Yet, the equilibrium towards the synthesis of formate may be disrupted at points when the NAD^+^/NADH balance must be restored. At this point, the pool of THF intermediates must be relieved from the MRO for the redox balance to be maintained and the cycle to continue (Fig 3).

Folate and its activated derivatives THF (H_4_folate, H_2_folate) require polyglutamylation to become carriers of the 1C moiety, and to maintain their compartmentalisation (Lawrence et al., 2014). Indeed, folylpolyglutamate synthase (FPGS; PaPyr1465) is encoded by the *P. pyriformis* genome and is likely dually localised between MRO and cytosol. The transfer of folate and its reduced derivatives between the mitochondrial and the cytosolic compartments has been thoroughly assessed. Isotope labelling experiments have been performed in a plethora of cell types and tissues, using inhibitors of the folate pathway, in aerobiosis/anaerobiosis, and with the aid of diversely labelled substrates (Cybulski and Fisher, 1981; Horne et al., 1992; Lichtenstein et al., 1969; Pasternack et al., 2002; Tan et al., 2020). Many of these results support the notion that certain THF species may be capable of passing from one cellular compartment to another in a carrier-mediated fashion, and the difference in the number of polyglutamylation chains seems to be the defining component of this behaviour. We hypothesise that some of the 4 MCF family proteins identified in *P. pyriformis* are likely involved in this process.

The presence of 1C metabolism known from aerobic mitochondria may not be limited to the MRO of *P. pyriformis*. Leger et al. (2017) predicted GCS and SHMT in the MROs of all free-living Fornicates and proposed that glycine degradation correlates with their lifestyle for an unclear reason. Moreover, GCS and SHMT were predicted to be localised in the MRO of other free-living Metamonada, such as *Barthelona sp*., *Anaeramoeba flamelloides* and *A. ignava* (Stairs et al., 2021; Yazaki et al., 2020) and free-living Archamoebae *Mastigamoeba balamuthii* and *Pelomyxa schiedtii* (Nývltová et al., 2015; Záhonová et al., 2022). Our data are consistent with these observations and provide a potential reason for maintaining GCS and SHMT in these organelles – production of folates to supply cytosolic 1C metabolism. Folate pathway enzymes have not been predicted in the MROs of other free-living metamonads, probably due to the limitations of *in-silico* methods thus were not queried as unambiguously mitochondrial proteins. Indeed, *P. pyriformis* FolD (PaPyr3253) and MTHFD (PaPyr5119), clearly classified as MRO proteins by LOPIT, were predicted as mitochondrial only by Cello and the fungal setting of MultiLoc2, whereas all other tools and settings considered them cytosolic. Finally, using *T. vaginalis* as a heterologous expression system provided false-negative results, as overexpressed FolD and MTHFD remained in the cytosol of *T. vaginalis*. In summary, we hypothesise that 1C metabolism remains an overlooked function of other MROs which cannot be easily revealed by *in silico* predictions but rather by proteomic studies similar to this work.

Several MRO proteins with weak homology to database representatives were present in the LOPIT dataset, and most of them enrich the inventory of membrane proteins responsible for organelle biogenesis. The protein translocation machinery of *P. pyriformis* appears to be reduced compared to *Trichomonas vaginalis* and anaeramoebids (Makki et al., 2019; Rada et al., 2011; Stairs et al., 2021), however, most of the core components were detected in the transcriptome or genome (Tom40, Sam50, Tim17, Tim23, Tim14, Cpn10, Cpn60, MPPαβ) (Zubáčová et al., 2013) and in our study, we further identified distant homologues of Tom22 (PaPyr4978), Tim44 (PaPyr7077), an extremely divergent Tim22 protein (PaPyr1418), and DnaK (PaPyr2959). The presence of a tail-anchored (TA) Rho GTPase Miro should be noted with particular interest. This protein is known from a wide spectrum of aerobic mitochondria, hypothesised to play roles in the regulation of mitochondrial morphology, transport along microtubules, and interactions with the ER (Yamaoka and Hara-Nishimura, 2014). Functional studies on *Dictyostelium discoideum*, however, showed that knock-out of Miro does not compromise the transport and distribution of mitochondria yet affects its metabolism (Vlahou et al., 2011). So far, the presence of Miro proteins in organisms with reduced mitochondria has been limited to *Blastocystis* (Gentekaki et al., 2017), so its distribution could be more widespread in MROs than previously considered. Finally, *P. pyriformis* apparently recruited the Pex3-Pex19 system, known to mediate sorting and insertion of peroxisomal membrane proteins, to import TA proteins to the MRO membrane. It was shown that Miro proteins can interact with Pex19 for its membrane insertion (Covill-Cooke et al., 2020). In *Saccharomyces cerevisiae*, Pex19 mediates the import of TA proteins into mitochondria (Cichocki et al., 2018) and Pex3 shows mitochondrial localisation at the early stage of *de novo* peroxisomal biogenesis in human fibroblasts (Sugiura et al., 2017). Our finding regarding the MRO localisation of Pex3 in absence of peroxisomes adds to the slowly growing body of evidence on the versatility of this system.

It is widely accepted that Fe-S cluster synthesis is a common function of mitochondria and their derivatives with the debatable exception of *Entamoeba histolytica* (Braymer et al., 2021; Maralikova et al., 2010; Mi-ichi et al., 2009). Replacement of the mitochondrial ISC pathway by the bacterial SUF system in Preaxostyla was likely a preadaptation for the loss of MRO in *Monocercomonoides exilis* (Karnkowska et al., 2016). The localisation of the SUF system in *P. pyriformis* had not been previously addressed experimentally, and here we provide clear evidence that it is indeed localised in the cytosol and does not contribute to the essential role of the MRO. Interestingly, two proteins assigned to the MRO proteome, hydrogenase maturases HydE and HydF, known to bear Fe-S clusters, raise the question of how these proteins are maturated. The presence of these proteins in the MRO is remarkable also in terms of their function, which involves the maturation of the H-cluster of HydA, alongside maturase HydG (Britt et al., 2020). Confusingly enough, the third maturase HydG was robustly classified as cytosolic together with four paralogues of HydA; one HydA remained unclassified and four were not detected by LOPIT. It is unclear, how the maturation of cytosolic HydA is achieved if the set of maturases is distributed across two compartments, however, a similar situation has been reported already in *T. vaginalis* and *G. intestinalis* (Lloyd et al., 2002; Smutná et al., 2022). The lack of evidence for the MRO localisation of HydA is consistent with the apparent lack of PFOs and ferredoxins in this compartment and indicates that the pathway of extended glycolysis has cytosolic localisation in *P. pyriformis*.

In conclusion, we demonstrate the feasibility of the LOPIT-DC method for non-model, non-axenic organisms, opening avenues to robustly determine the localisation of hundreds of proteins simultaneously. Via this approach, we identified several new components of *P. pyriformis* MRO, including its protein import machinery and additional proteins partaking in its biogenesis. From the reconstructed biochemical map, we infer that the organelle provides 1C units to the cytosol to supply the methionine cycle. This points to folate metabolism, known from aerobic mitochondria, as another basic mitochondrial function until now overlooked in MROs but potentially very common among free-living anaerobes, most of which had already been shown to contain GCS and SHMT without satisfactory functional explanation. In the particular case of *P. pyriformis*, production of THF species and formate is probably the only reason why the cell maintains the organelle even after Fe-S cluster synthesis relocation to the cytosol.

## Acknowledgements

This project has received funding from the European Research Council (ERC) under the European Union’s Horizon 2020 research and innovation programme (grant agreement No 771592) and the Grant Agency of Charles University (Project No. 1162320). We would like to thank to Jan Tachezy for providing anti-malic antibody, Kristina Záhonová for providing Pex proteins alignments, Konstantin Barylyuk for useful comments regarding data processing and Jan Pyrih for discussions on 1C metabolism. We acknowledge Imaging Methods Core Facility at BIOCEV, institution supported by the MEYS CR (Large RI Project LM2018129 Czech-BioImaging) and ERDF (project No. CZ.02.1.01/0.0/0.0/18_046/0016045) for their support with obtaining imaging data presented in this paper. Computational resources were supplied by the project “e-Infrastruktura CZ” (e-INFRA CZ LM2018140) supported by the MEYS CR.

## Author contributions

V.H., conceived the study. P.P.D and V.H supervised the project. J.Z., performed DC-LOPIT and analyzed data with S.C.T., K.H., performed mass spectrometry analysis; Z.F., phylogenetic analyses and evaluation of targeting predictors; Z.V., D.Z., and J.Z., experiments with heterologous expression system. Z.F., and J.Z., prepared the figures. V.H., J.Z., Z.F., and P.P.D wrote the manuscript. All authors read and approved the manuscript.

## Declaration of interests

The authors declare no competing interests.

## Supplementary figure legends

**Figure S1. Electron microscopy of first six sub-cellular fractions**. Transmission electron microscopy showed that MROs remained intact during cells disruption and fractionation. Fractions of sub-cellular compartments were collected by differential centrifugation at different speeds A) 1,000 × *g*, B) 3,000 × *g*, C) 5,000 × *g*, D) 9,000 × *g*, E) 12,000 × *g*, F) 15,000 × *g* (scale bar, 2 μm) G) Enlarged intact MRO; arrows indicate double membrane.

**Figure S2. Sub-compartments resolution of LOPIT-DC quantified by Qsep tool**. A) Heatmap displays average normalised ratio of distances between clusters to within cluster. B) Boxplot summarizes Qsep average normalised distances. The dashed vertical line represents the overall median value of normalised pairwise distances.

**Figure S3. PCA projection after SVM classification with highlighted proteins of interest and distribution profile of FPGS**. A) PCA projection after SVM classification with highlighted HydA, hydrogenase maturases, PFOs, ferredoxins, FPGS and proteins from the SUF pathway. B) distribution profile of FPGS across then labelled fractions.

**Figure S4. Phylogenetic analyses of hypothetical MRO proteins of *P. pyriformis***. For three hypothetical proteins for which we could retrieve homological sequences in eukaryotes, we constructed ML phylogenies using the IQ-TREE’s Posterior Mean Site Frequency model and 1,000 ultra-fast bootstrap replicates (see Methods). The three hypothetical proteins, highlighted by black background, were identied as TRIC ion channel (PaPyr804), Tim44 (PaPyr7077), and peroxin-3 (PaPyr5495). Support values are shown where ≥ 85. Albeit the positions of *P. pyriformis* sequences are not well-supported, their homology with the respective protein families is unequivocal. Along with peroxin-3, the phylogeny of its interacting partner peroxin-19 is shown. In the lower panel, the alignment of selected Pex19 homologs is shown with the Pex3-interacting domain highlighted in red. Although this is not recognized by the alignment algorithm, this domain is characteristic by a sequence of hydrophobic and charged amino acids. The core peroxin-19 domain is located downstream. In case of PaPyr8690, this domain is not identified by HHpred or Pfam and was inferred by homology with the sequence from *Trimastix marina* (shown in opaque colors).

**Figure S5. Localisation of *P. pyriformis* proteins using *Trichomonas vaginalis* as a heterologous expression system**. *P. pyriformis* proteins C-terminally HA-tagged were expressed in *T. vaginalis* and visualised by immunofluorescence microscopy. Anti-HA antibody (cyan) detected overexpressed proteins; malic enzyme (ME, hydrogenosomal marker) was detected with polyclonal anti-ME antibody (magenta). DAPI was used as nuclear stain (blue). The scale bar represents 5 μm. HydA_2, hydrogenase (PaPyr1278); HydA_4, hydrogenase (PaPyr6388); HydA_6, hydrogenase (PaPyr8158); HydE, hydrogenase maturase E (PaPyr4383); HydF, hydrogenase maturase F (PaPyr9283); HydG, hydrogenase maturase G (PaPyr1252); PFO_2 pyruvate:ferredoxin oxidoreductase (PaPyr12211); PFO_4, pyruvate:ferredoxin oxidoreductase (PaPyr7937_PaPyr7938); FolD, methylenetetrahydrofolate dehydrogenase/methenyl tetrahydrofolate cyclohydrolase (PaPyr3253); MTHFD, formate-tetrahydrofolate ligase (PaPyr5119); SufC (PaPyr9323); VDAC, voltage-dependent anion channel (PaPyr2234); Tom22 (PaPyr4978); Pex3, peroxisomal biogenesis factor 3 (PaPyr5495).

**Table S1. Non-normalised abundance of TMT-labelled peptides and proteins metadata**.

**Table S2. Normalised abundance of TMT-labelled peptides and outcomes of classification methods**.

**Table S3. Calculated net charge of the first 10, 20 and 30 amino acids of MRO proteins**.

**Table S4. Evaluation of targeting predictors**.

**Table S5. Sequences of primers used for heterologous expression system**.

## Materials and Methods

### Cell cultivation

*Paratrimastix pyriformis* (strain RCP-MX, ATCC 50935) was grown at room temperature in a polyxenic culture in Sonneborn’s Paramecium medium ATCC 802 (Sonneborn, 1950) supplemented with 1% (v/v) of Luria-Bertani (LB) broth (Bertani, 1951).

*Trichomonas vaginalis* (strain T1) was maintained at 37°C in tryptone-yeast extract-maltose medium (TYM, pH 6.3) supplemented with 10% (v/v) heat-inactivated horse serum (Diamond, 1957).

### Subcellular fractionation by differential centrifugation

Prior to fractionation, 6-7 L of *P. pyriformis* culture on exponential phase of growth was filtered through filter paper (80 g/m^2^) to remove bacterial aggregates, and subsequently sieved through a 3 μm polycarbonate filter (Nuclepore Track-Etch Membrane,Whatman) followed by two washing steps with 10% (v/v) LB. *P. pyriformis* cells were collected into a 50 mL tube and harvested by centrifugation at 1000 × *g* for 10 min at room temperature. The cell pellet (approximately 2 × 10^8^ cells) was resuspended in 2 mL of pre-cooled lysis buffer (10 mM HEPES pH 7.4, 25 mM sucrose, 2 mM magnesium acetate, 2 mM EDTA) supplemented with cOmplete EDTA-free protease inhibitors (Roche). Subcellular fractionation by differential centrifugation was performed as described in (Geladaki et al., 2019) with slight modifications. Briefly, cells were disrupted by three rounds of sonication of 10 s with pulse 1 s ON / 1 s OFF at amplitude 20%, using a standard 1/8” diameter probe (Qsonica). After each round, cells were examined under the microscope. Three rounds were sufficient to break 80-90% of cells. The sonicate was centrifuged twice at 200 × *g* at 4 °C for 10 min to remove unbroken cells. Nine fractions pelleted at different centrifugation speeds were collected and the final supernatant after 120,000 × *g* represents the cytosol-enriched fraction. Collected pellets were resuspended in 50-100 μL of membrane solubilising buffer [50 mM HEPES pH 8.5, 8 M urea, 0.2 % (w/v) SDS]. Protein concentration was measured by BCA Protein Assay Kit (Sigma-Aldrich) according to manufacturer’s instructions.

### Protein digestion and TMT labelling

A total of 60 μg of protein from each fraction was bound to magnetic HILIC beads according to the SP3 protocol published by (Hughes et al., 2018). After washing out the buffer with 80% (v/v) EtOH, the protein-bead mixture was resuspended in 100 mM triethylammonium bicarbonate (TEAB), disulphide bridges were reduced with 10 mM Tris(2-carboxyethyl)phosphine (TCEP) and cysteines were modified with 50 mM chloroacetamide (one step reaction 30 min at 60 °C). To digest proteins, 2 μg of trypsin was added to the reaction and incubated overnight at 37 °C. The next day, samples were centrifugated and the collected supernatants were acidified with trifluoroacetic acid (TFA) to a final concentration 1% (v/v). At this step, samples were divided into technical triplicates. 20 μg of peptides were desalted on a homemade desalting column (Kulak et al., 2014), reconstituted in 17μL 100mM TEAB, and labelled with TMT 10plex isobaric tagging reagent (Thermo Fisher Scientific). 0.8 mg of TMT 10plex reagents were resuspended in 42 μL of anhydrous acetonitrile (ACN). Then 6.4 μL of the TMT reagent was added to each fraction sample and incubated for one hour at room temperature. The reaction was stopped by the addition of 1.5 μL of 5% (v/v) hydroxylamine. Subsequently, the respective indicated fractions were mixed. After evaporation, the sample was desalted on a Opti-trap cartridge C18 column (Optimize Technologies).

The resulting mixture was fractionated on a reverse phase at high pH, flow 2 μL/min, YMC column (300 mm, 0.3 mm, 1.9 μm), linear 60 min gradient [buffer A – 20 mM ammonium formate, 2% (v/v) ACN; buffer B – 20 mM ammonium formate, 80% (v/v) ACN] from 1% B to 60% B (Kulak et al., 2017). Fractions were collected for 64 minutes, and these were combined into 8 resulting pooled fractions by the farthest neighbour method described in (Wang et al., 2011)

The fractionated peptides were resuspended in 1% (v/v) TFA + 2% ACN and roughly 1.5 μg were injected onto nanoHPLC Dionex UltiMate 3000RS (Thermo Fisher Scientific) in combination with Thermo Orbitrap Fusion. Thermo PepMap Trap column (pn: 160454) was used for peptides preconcentration and main separation was done on 50 cm EASY Spray Column Thermo (pn: ES903). Each fraction was separated on a 180 min gradient from 2% A to 35% B [A – 0.1% (v/v) formic acid, B – 99.9% ACN, 0.1% (v/v) formic acid]. Cycle time was set to 4 s. The MS^2^ spectra for identification were measured in an ion trap with CID fragmentation, 60 ms maximum injection time, and the quantification MS^3^ spectra were measured in an Orbitrap with HCD fragmentation using the SPS function, 118 ms maximum injection time, 10 precursors for synchronous precursors selection feature (Le et al., 2020).

### Raw LC-MS data processing

Raw data from each biological replicate were searched with Proteome Discoverer 2.4 (Thermo Fisher Scientific) with Sequest search engine. Method was based on predefined workflow for SPS MS3 isobaric quantification and was used with batch specific correction parameters for TMT 10plex label. Database of *Paratrimastix pyriformis* (13,533 entries) was used to search along with common contaminants database and databases of selected bacterial taxa known to be co-cultivated with *P. pyriformis* (ASM2556v1 *Enterobacter cloacae*, ASM63591v2 *Brevundimonas naejangsanensis*, ASM64851v1 *Citrobaster freundii*, ASM76115v1 *Pseudomonas rhizosphaerae*, ASM383324v1 *Paenibacillus xylanexedens*, ASM385183v1 *Pseudomonas chlororaphis* and GCF_900105615.1 *Clostridium sphenoides*). Dynamic modifications were set as follows: Oxidation / +15.995 Da (M). Protein N-Terminal: Acetyl / +42.011 Da, Met-loss / -131.040 Da (M), Met-loss+Acetyl / -89.030 Da (M). Static Modifications: Carbamidomethyl / +57.021 Da (C), TMT6plex / +229.163 Da (K), Peptide N-Terminus: TMT6plex / +229.163 Da. To assess FDR, percolator was used. For reporter ions detection 20 pmm tolerance was set with most confident centroid peak integration.

### LOPIT-DC data processing

Data were processed in R (version 4.0.4) using pRoloc package (version 1.30.0) (Gatto and Christoforou, 2014) according to the published Bioconductor workflows (Breckels et al., 2018; Crook et al., 2019). Briefly, experimental replicates were pre-processed to allow decontamination and normalisation. Bacterial and other contaminants were removed. Then, average reporter intensity was calculated for each feature based on three technical replicates of the LC-MS measurement. All proteins that contained one or more missing values across ten labelled fractions were removed from the dataset by filterNA function. Normalisation of protein intensity-interval was performed by the sum method (each reporter intensity was divided by the sum of the reporter ions intensities). After normalisation, four independent LOPIT-DC experiments were merged into one dataset containing 4,541 proteins common for all four replicates.

To search for the subcellular compartments, we used the HDBSCAN unsupervised clustering (Campello et al., 2013) available in Python, and semi-supervised phenotype discovery algorithm (phenoDisco) (Breckels et al., 2013) available in the pRoloc package of R. The following parameters were applied for HDBSCAN: min_cluster_size=10, min_samples=8, cluster_selection_method=leaf’. Phenotype discovery was run several times with the group size (GS, minimum number of proteins per group) set to 10, 20 or 30. The content of newly found clusters by either HDBSCAN or phenoDisco were carefully evaluated. Finally, by using these two methods as well as gene annotation, conserved domains, previous studies, and literature, a list of 226 manually curated proteins defining 11 cell sub-compartments (cytosol, proteasome, ribosome, nucleus, nucleus-chromatin, ER, Golgi, endomembrane vesicles, plasma membrane, MRO, food vacuole) was prepared. Different populations of endosomes were merged to form a single class, and the same was performed for 40S/60S ribosomal subunits.

To assign proteins with unknown localisation into subcellular clusters two supervised machine learning approaches were used: support vector machine (SVM) and t-augmented Gaussian mixture (TAGM) model (Crook et al., 2018) both available in pRoloc package. In the case of SVM the parameters were optimised as described (Breckels et al., 2018). The best sigma and cost parameters were chosen (sigma = 0.1, cost = 8) based on F1 score. For most clusters, median was used to set up the cut-off scores. Threshold scores were as follows: Cytosol = 0.55, Proteasome = 0.53, Ribosome = 0.63, Nucleus = 0.5, Nucleus-chromatin = 0.42, ER = 0.63, Golgi = 0.4, Endomembrane vesicles = 0.45, Plasma membrane = 0.53, MRO = 0.55, Food vacuole = 0.3. Any prediction of protein localisation with a score below these thresholds was labelled as unknown. TAGM MAP, the other used classification method, assigns proteins into sub-compartments based on a probabilistic approach (Crook et al., 2018). Uniform localisation threshold set to 99% probability was applied for this method. To reduce dimensions and visualise the data, principal component analysis (PCA) was used.

### RNA extraction and cDNA synthesis

Total RNA was extracted from 2 L of *P. pyriformis* culture using TRI Reagent (Sigma-Aldrich) according to the manufacturer’s instructions. To prepare cDNA, mRNA was isolated from total RNA using Dynabeads mRNA Purification Kit (Invitrogen), followed by cDNA synthesis using SMARTer PCR cDNA synthesis Kit (Takara Bio).

### Localisation of *P. pyriformis* proteins by heterologous expression system

Selected genes of *P. pyriformis* were amplified using PrimeSTAR MAX DNA Polymerase (Takara Bio) from cDNA and cloned into the TagVag2 vector (Hrdy et al., 2004) via restriction endonucleases or Gibson assembly. Sequences of primers used for cloning are summarized in Table S5. Electroporation of *Trichomonas vaginalis* was performed as described (Novák et al., 2016). Four hours after electroporation, G418 (200 μg/mL) was added to medium. After 3-6 passages expression of *P. pyriformis* genes in *T. vaginalis* cells was examined by immunoblot and immunofluorescence microscopy.

### Immunofluorescence microscopy

Slides for immunostaining were prepared as described previously in (Dawson et al., 2008) with slight modifications. *T. vaginalis* cells were fixed with 4% (v/v) formaldehyde for 30 min at room temperature. Fixed cells were spread on coverslips coated with 0.1% (v/v) poly-L-lysine solution then left 30 min to adhere and air-dried.

*T. vaginalis* cells were stained with anti-HA rat monoclonal antibody (Roche) at dilution 1:500 to examine subcellular localisation of overexpressed *P. pyriformi*s genes. Hydrogenosomes were stained with rabbit anti-malic enzyme antibody (kindly provided by prof. Jan Tachezy) at a dilution of 1:500. Secondary antibodies Alexa488 Goat anti-Rat (Invitrogen) and Alexa594 Goat anti-Rabbit (Invitrogen) were used at dilution 1:1000. All antibodies were diluted in PEMBALG buffer [100 mM PIPES, pH 6.9, 1 mM EGTA, 0.1 mM MgSO_4_, 100 mM lysine, and 0.5% (w/v) cold water fish skin gelatine]. Coverslips were mounted to the slides using VECTASHIELD Mounting Medium with DAPI (Vector Laboratories).

Images of prepared slides were captured using a Leica SP8 confocal microscope. Deconvolution of images were performed using Huygens Professional (version 21.10). Deconvolved images were further processed using Fiji software (version 1.53f51) (Schindelin et al., 2012).

### Immunoblotting

To confirm expression and size of *P. pyriformis* proteins expressed in *Trichomonas vaginalis*, immunoblotting was performed. Approximately 5×10^6^ of *T. vaginalis* cells were harvested by centrifugation at 1000 × *g* for 10 min to prepare a whole cell lysate. Pelleted cells were resuspended in 50 μL of 1 × SDS-PAGE sample buffer and denatured for 5 min at 95°C. Ten μL of each sample was resolved by SDS-PAGE using 12% (v/v) polyacrylamide gel. Samples were electrophoretically transferred onto PVDF membranes (Amersham). Non-specific binding was blocked by incubation of the membranes in 5% (w/v) non-fat milk in TBST [Tris-buffered saline, 0.1% (v/v) Tween 20]. Membranes were incubated overnight at 4°C with primary anti-HA antibody, washed 3×10 min with TBST and incubated with a secondary antibody conjugated to HRP. After 1 h of incubation, membranes were washed 3×10 min with TBST. To visualise protein bands via chemiluminescence, Clarity ECL Western Blotting Substrates (Bio-Rad) was applied to the PVDF membrane and detected on an Amersham Imager 600.

### Electron microscopy

Approximately 3 L of culture were filtered and fractionated as described above. Pellets of the first six collected fractions were resuspended in 1 mL of fixation buffer containing 200 mM sodium cacodylate (pH 7.4) and 2.5% (v/v) glutaraldehyde. After 3 h of incubation at room temperature, samples were washed twice by 100 mM sodium cacodylate. Next, post-fixation with 2% (v/v) OsO_4_ in 100mM sodium cacodylate (pH 7.4) buffer was carried out on ice for 1 hour. Dehydration of fixed subcellular fractions was performed through an ethanol series (35, 50, 70, 80, 95, 100%) and then ethanol was replaced with 100% acetone. Infiltration with a mixture of Epon resin and 100% acetone at a ratio 1:2, 1:1, 2:1 (one hour each) was followed by overnight incubation with absolute Epon resin. After polymerisation of the resin (65°C, 48 h) ultrathin sections were prepared using a Reichert-Jung Ultracut E ultramicrotome and collected onto carbon-and-Formvar coated copper grids. Ultrathin sections were post-contrasted with 4% (w/v) uranyl acetate and 2% (w/v) lead citrate. Prepared sections were examined JEOL 1011 transmission electron microscope.

### Phylogenetic and in silico structural analyses

To clarify the identity of the hypothetical proteins of the *P. pyriformis* MRO, we employed a combination of phylogenetic reconstruction and in silico structure prediction. For three proteins (PaPyr5495, PaPyr7077 and PaPyr804) we could retrieve multiple BLAST hits, based on which we created HMMER v3.3 (www.hmmer.org) profiles for a more comprehensive search of homologues in the NCBI nonredundant protein (nr) database and EukProt (Richter et al., 2020). Regions flanking the domains of interest as determined by InterProScan v5.52-86.0 (Jones et al., 2014) were manually removed in Geneious Prime v2020.1.2 (Kearse et al., 2012) to allow more reliable sequence alignments. The resulting datasets were aligned by MAFFT v7.453 (Katoh and Standley, 2013) using the L-INS-i refinement and a maximum of 1,000 iterations, followed by manual trimming; alignments are attached as Supplementary Material. ML trees were inferred by IQ-TREE v 1.6.12 using the Posterior Mean Site Frequency (PMSF) empirical model with ultrafast bootstrapping strategy (1,000 replicates) and a LG4X guide tree (Wang et al., 2018, 2019). Obvious contaminants were manually pruned based on intermediary phylogenies as well as functional domain predictions by InterProScan. For PaPyr577, a single homologue was found in *Trimastix marina* (AC: Trimastix_PCT|3971) and the resulting HMMer profile did not recover more homologues in EukProt. To find Pex19 (PaPyr8690), and the members of Tim17/22/23 protein family (PaPyr5590, PaPyr9, PaPyr1418), we used previously constructed alignments kindly provided by Kristína Záhonová and Vojtěch Žárský (Žárský and Doležal, 2016). In addition, we analysed the phylogenetic relationships of enzymes of the cytosolic and MRO folate pathways. Their phylogenetic analysis was performed as above.

For the hypothetical proteins lacking any sequence homologues and thus phylogenetic support, SWISS-MODEL (Waterhouse et al., 2018) and PHYRE2 (Kelley et al., 2015) structure-based searches were performed (last online access in October 2021) to identify putative functions.

### Testing of MRO targeting prediction

For performance comparison of targeting predictors, 30 complete, reliably predicted proteins from the MRO were selected along with 30 complete, randomly chosen, and reliably predicted proteins from each of the remaining compartments. These were analysed by the following algorithms: Cello v2.5 in eukaryote mode (last accessed 5 February 2022) (Yu et al., 2006), iPSORT in non-plant mode (Bannai et al., 2002), MitoFates v1.1 in Metazoa mode (Fukasawa et al., 2015), MultiLoc2 in animal and fungal modes (Blum et al., 2009), NommPred v0.2 in MRO mode (Kume et al., 2018) and TargetP v2.0 in non-plant mode (Armenteros et al., 2019). From tabulated results, sensitivity and specificity were calculated using formulae

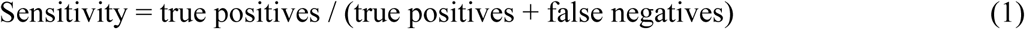

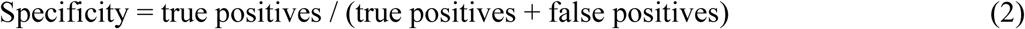

Other than default thresholds were also considered when optimising these statistics. Notably, ribosomal proteins were more frequently predicted as mitochondrial by Cello and their omission from the tested sequences led to increased specificity (0.57 vs. 0.41).

## Resource availability

### Lead contact

Further information and requests for resources and reagents should be directed to and will be fulfilled by the lead contact, Vladimír Hampl.

### Materials availability

Plasmids and T. vaginalis cell lines generated in this study are available from the lead contact upon request.

### Data and code availability

The predicted proteome of *Paratrimastix pyriformis* is available at https://doi.org/10.5281/zenodo.6405158. The mass-spectrometry-based proteomics data have been deposited to the ProteomeXchange Consortium (Deutsch et al., 2017) via the PRIDE (Perez-Riverol et al., 2019) partner repository.

## Notes

https://doi.org/10.5281/zenodo.6405158

### Competing Interest Statement

The authors have declared no competing interest.

