## Supplementary figures and images for "Reduced mitochondria provide an essential function for the cytosolic methionine cycle"

### Figure S1

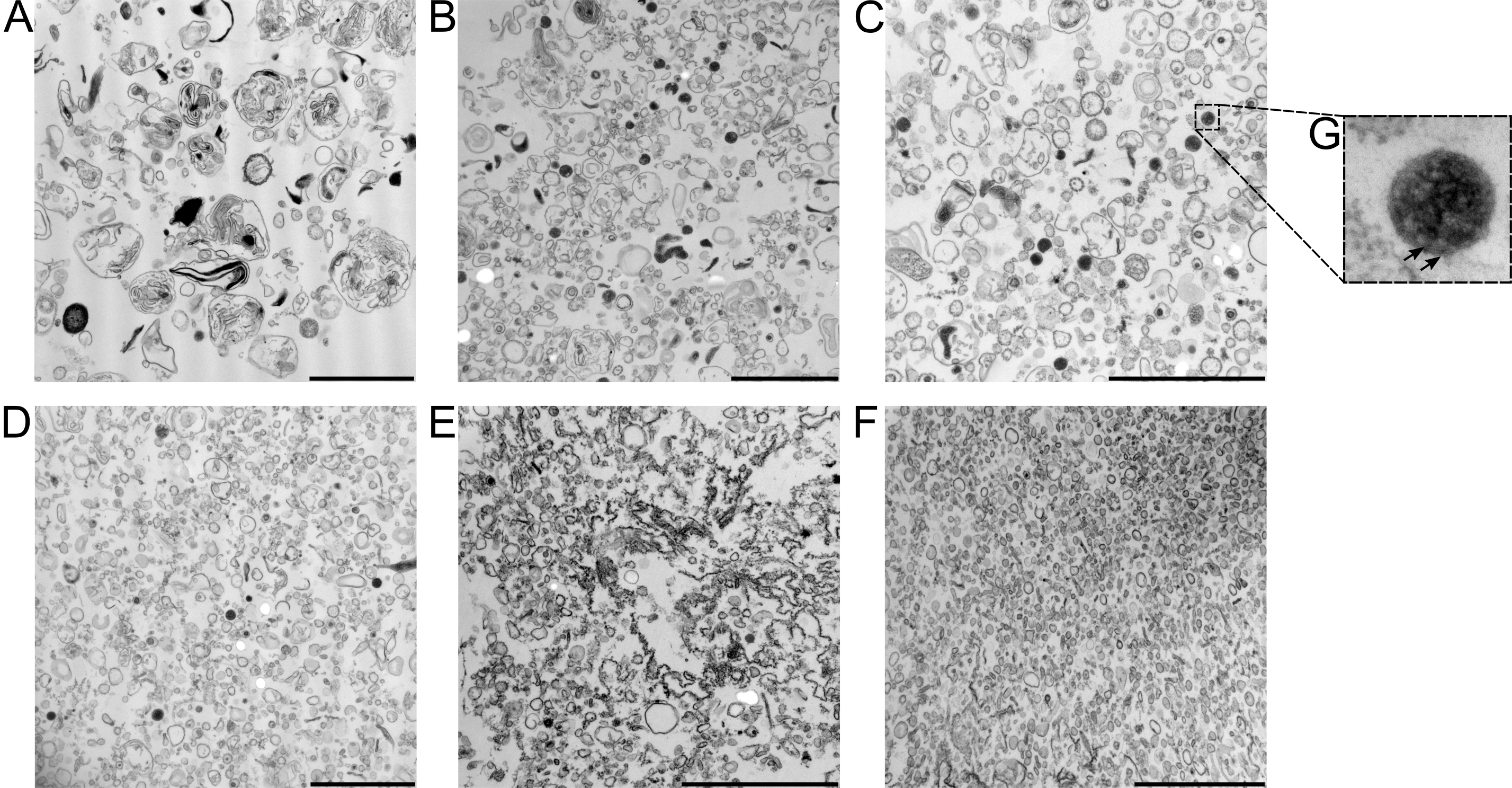

### Figure S2

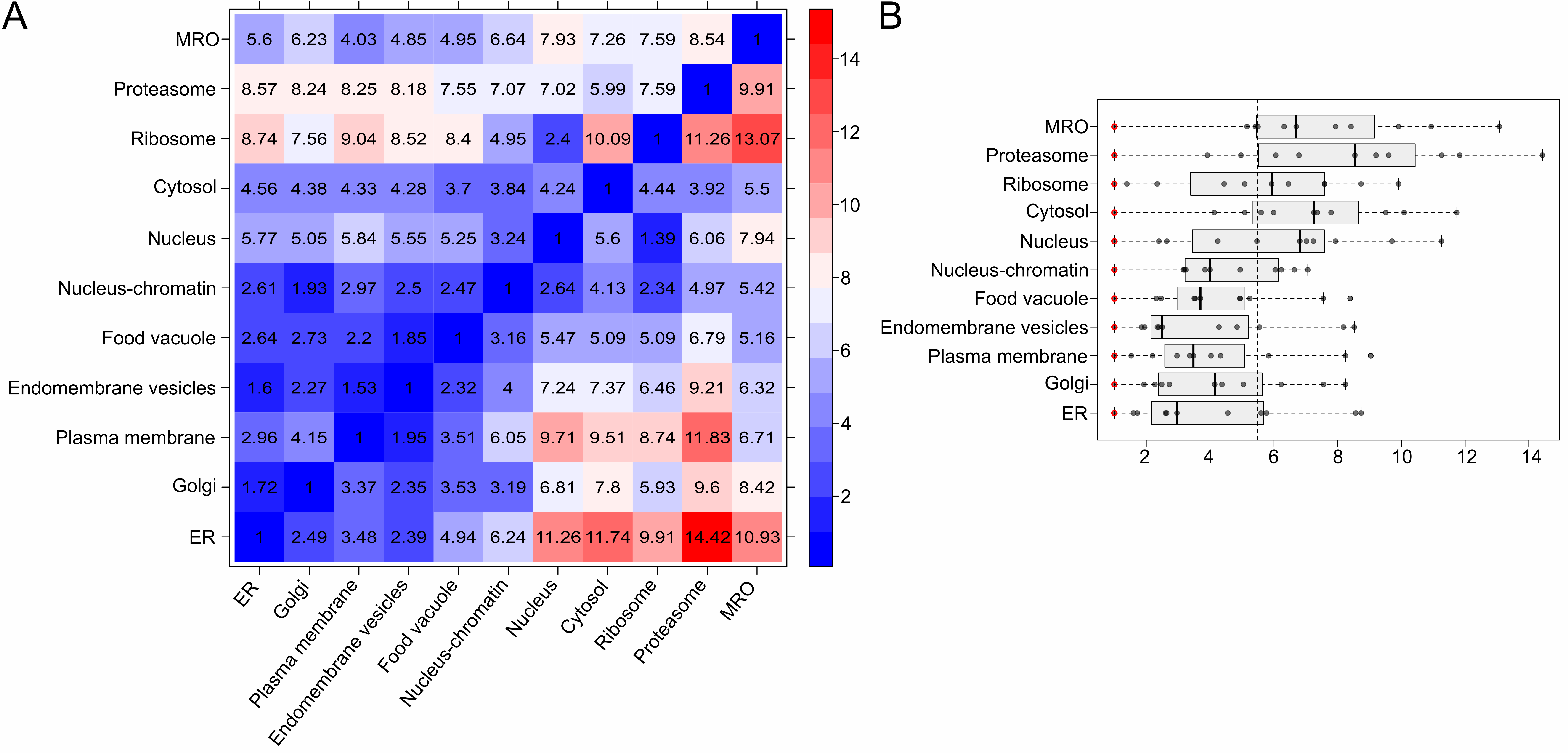

### Figure S3

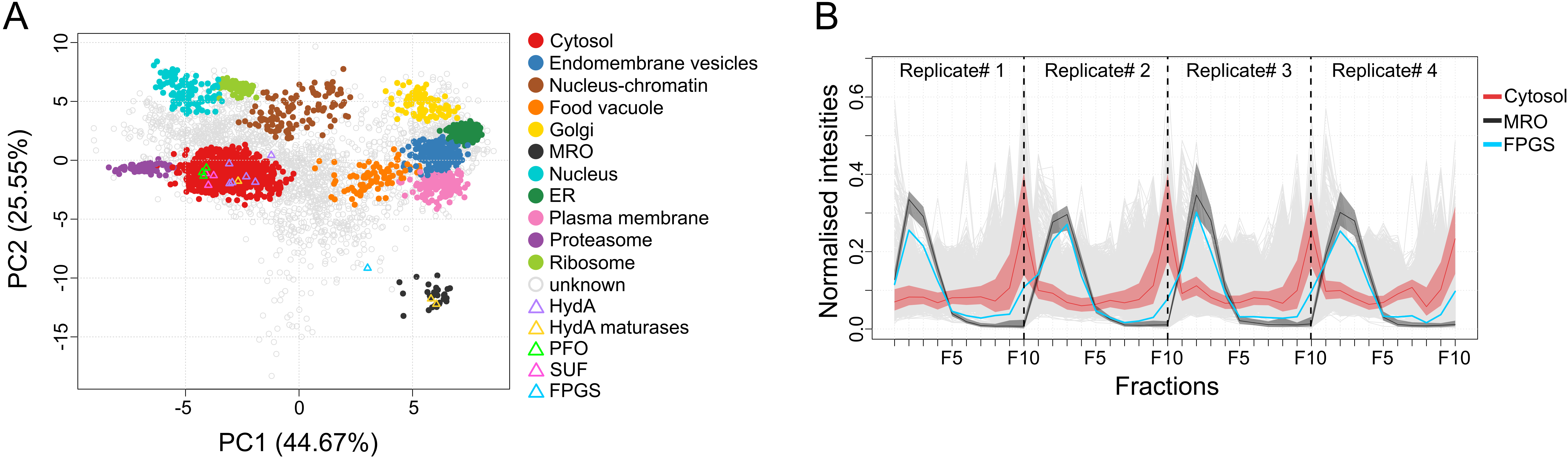

### Figure S5

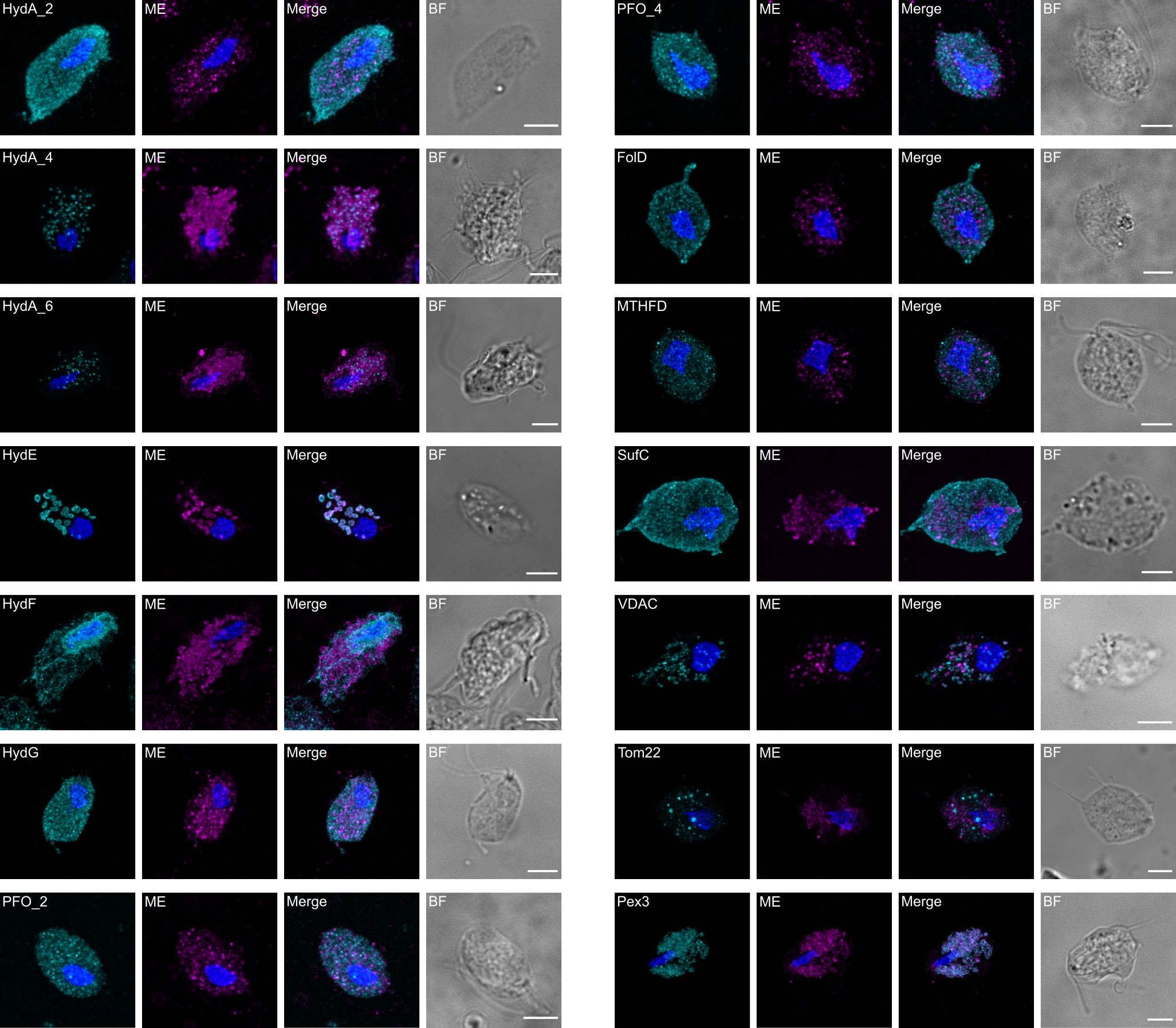
