## Supplementary material for "Reduced mitochondria provide an essential function for the cytosolic methionine cycle": Figure S4

**SUPPLEMENTARY FIGURE S4: Phylogenetic analyses of hypothetical MRO proteins of *P. pyriformis*.** For three hypothetical proteins for which we could retrieve homological sequences in eukaryotes, we constructed ML phylogenies using the IQ-TREE's Posterior Mean Site Frequency model and 1,000 ultra-fast bootstrap replicates (see Methods). The three hypothetical proteins, highlighted by black background, were identified as TRIC ion channel (PaPyr804), Tim44 (PaPyr7077), and peroxin-3 (PaPyr5495). Support values are shown where ≥ 85. Albeit the positions of *P. pyriformis* sequences are not well-supported, their homology with the respective protein families is unequivocal.
Along with peroxin-3, the phylogeny of its interacting partner peroxin-19 is shown. In the lower panel, the alignment of selected Pex19 homologs is shown with the Pex3-interacting domain highlighted in red. Although this is not recognized by the alignment algorithm, this domain is characteristic by a sequence of hydrophobic and charged amino acids. The core peroxin-19 domain is located downstream. In case of PaPyr8690, this domain is not identified by HHpred or Pfam and was inferred by homology with the sequence from *Trimastix marina* (shown in opaque colors).

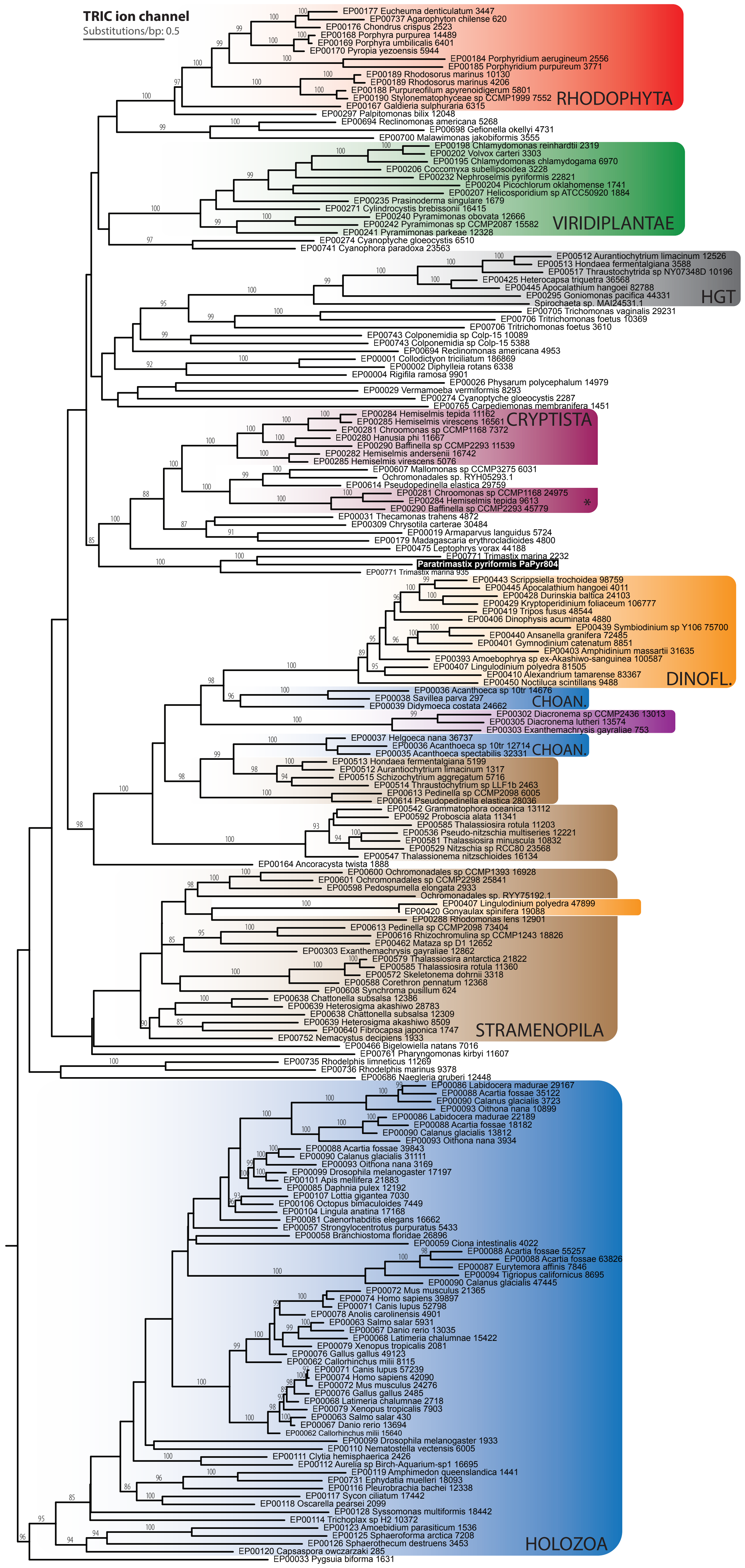

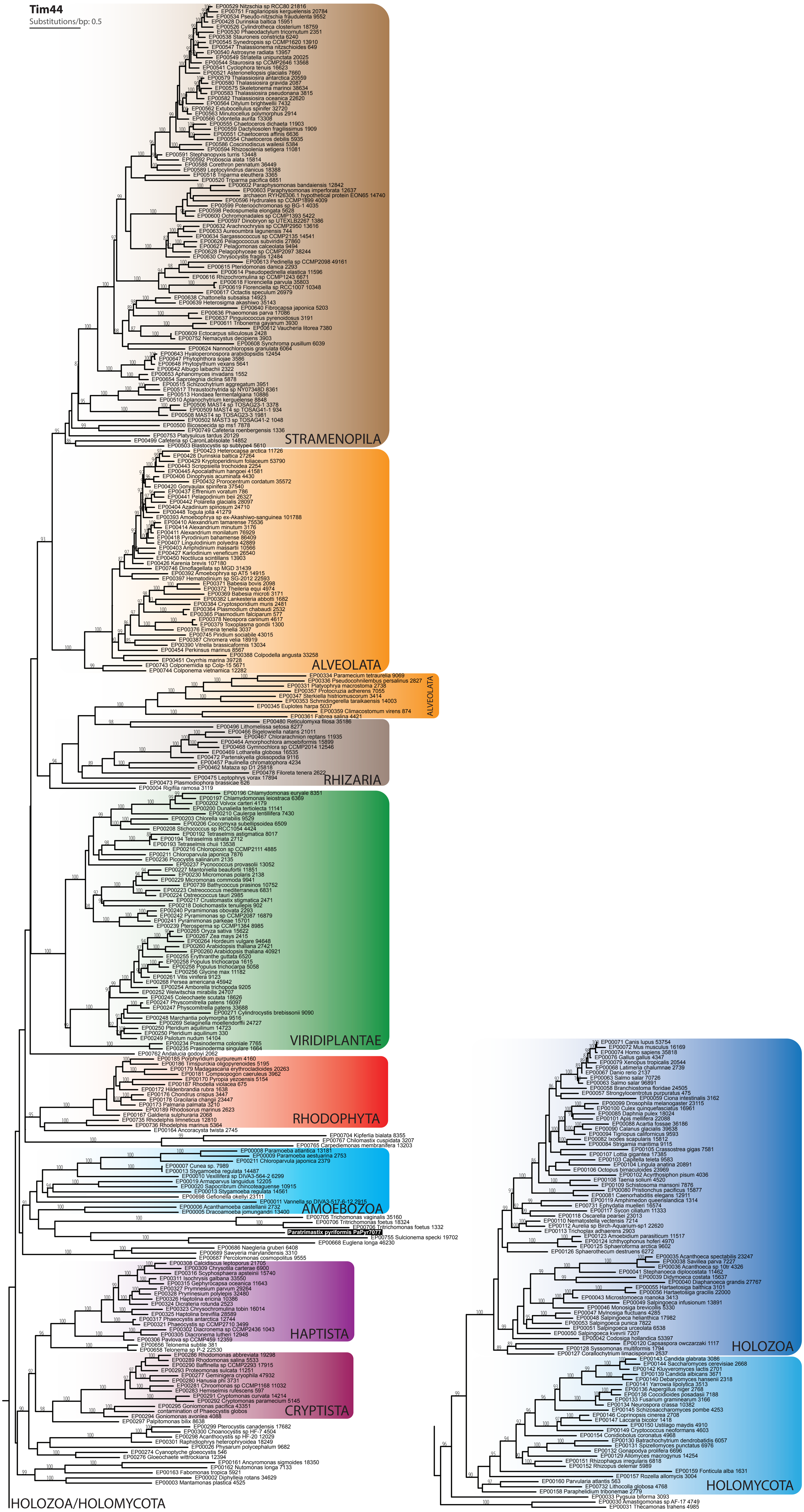

Tim44
Substitutions/bp: 0.5
STRAMENOPILA
ALVEOLATA
ALVEOLATA
RHIZARIA
VIRIDIPLANTAE
RHODOPHYTA
AMOEBOZOA
Paratrimastix pyriformis PaPyr7077
HAPTISTA
CRYPTISTA
HOLOZOA/HOLOMYCOTA
HOLOZOA
HOLOMYCOTA

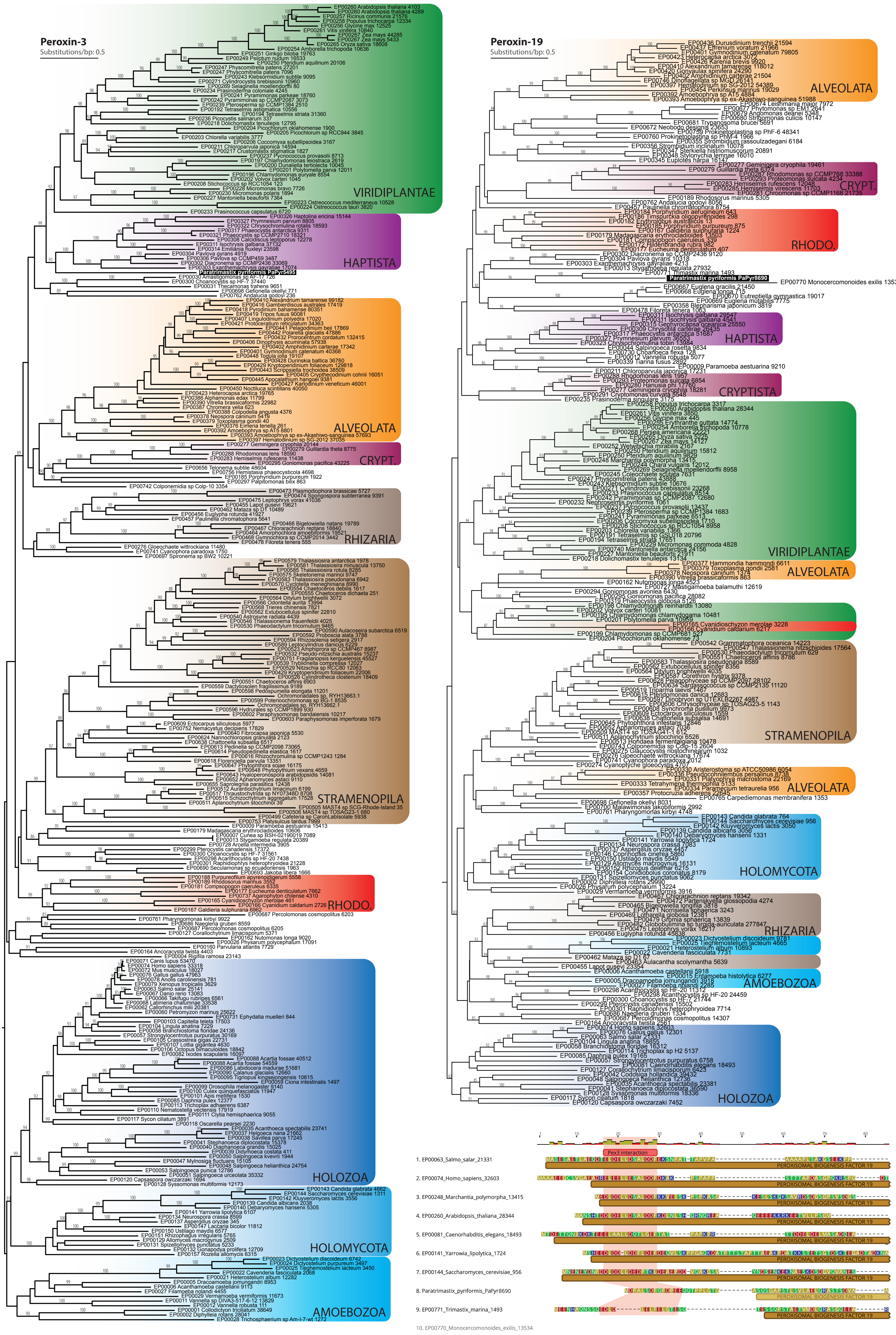

Peroxin-3
Substitutions/bp: 0.5
VIRIDIPLANTAE
HAPTISTA
Paratrimastix pyriformis PaPyr5495
ALVEOLATA
CRYPT.
RHIZARIA
STRAMENOPILA
RHODO.
HOLOZOA
HOLOMYCOTA
AMOEBOZOA
Peroxin-19
Substitutions/bp: 0.5
ALVEOLATA
CRYPT.
RHODO.
Paratrimastix pyriformis PaPyr8690
EP00770 Monocercomonoides exilis 13534
HAPTISTA
CRYPTISTA
VIRIDIPLANTAE
ALVEOLATA
STRAMENOPILA
ALVEOLATA
HOLOMYCOTA
RHIZARIA
AMOEBOZOA
HOLOZOA
Pex3 interaction
1. EP00063_Salmo_salar_21331
2. EP00074_Homo_sapiens_32603
3. EP00248_Marchantia_polymorpha_13415
4. EP00260_Arabidopsis_thaliana_28344
5. EP00081_Caenorhabditis_elegans_18493
6. EP00141_Yarrowia_lipolytica_1724
7. EP00144_Saccharomyces_cerevisiae_956
8. Paratrimastix_pyriformis_PaPyr8690
9. EP00771_Trimastix_marina_1493
10. EP00770_Monocercomonoides_exilis_13534
PEROXISOMAL BIOGENESIS FACTOR 19
